# Organellular imaging *in vivo* reveals a depletion of endoplasmic reticular calcium during post-ictal cortical spreading depolarization

**DOI:** 10.1101/2024.09.21.614252

**Authors:** Matthew A. Stern, Eric R. Cole, Claire-Anne Gutekunst, Jenny J. Yang, Ken Berglund, Robert E. Gross

**Affiliations:** Department of Neurosurgery, Emory University School of Medicine, Atlanta, GA, United States; Coulter Department of Biomedical Engineering, Emory University and Georgia Institute of Technology, Atlanta, GA, United States; Department of Chemistry, Center for Diagnostics and Therapeutics, Advanced Translational Imaging Facility, Georgia State University, Atlanta, GA, United States

## Abstract

During cortical spreading depolarization (CSD), neurons exhibit a dramatic increase in cytosolic calcium, which may be integral to CSD-mediated seizure termination. This calcium increase greatly exceeds that during seizures, suggesting the calcium source may not be solely extracellular. Thus, we sought to determine if the endoplasmic reticulum (ER), the largest intracellular calcium store, is involved. We developed a two-photon calcium imaging paradigm to simultaneously record the cytosol and ER during seizures in awake mice. Paired with direct current recording, we reveal that CSD can manifest as a slow post-ictal cytosolic calcium wave with a concomitant depletion of ER calcium that is spatiotemporally consistent with a calcium-induced calcium release. Importantly, we observed both naturally occurring and electrically induced CSD suppressed post-ictal epileptiform activity. Collectively, this work links ER dynamics to CSD, which serves as an innate process for seizure suppression and a potential mechanism underlying therapeutic electrical stimulation for epilepsy.

## Introduction

The endoplasmic reticulum (ER), a major calcium reservoir of the cell, is critically involved in essential physiological processes. The perturbation of ER homeostatic regulation in turn has serious pathophysiologic implications^1^. This is especially the case for the nervous system, where intra- and inter-cellular communication is tightly regulated through the ER calcium, including synaptic transmission, transcriptional regulation and plasticity^2–4^. Deciphering calcium signaling dynamics at a spatiotemporal level during aberrant activity could inform our understanding of disease mechanisms and their downstream sequelae. Calcium imaging of the ER has been enabled through the development of various ER targeted dyes and genetically encoded calcium indicators^5, 6^, but these have hitherto not been applied *in vivo* to vertebrates. Thus, we devised an approach for concurrent *in vivo* calcium imaging of the cytosol and ER. We then applied this approach to elucidate the difference between two related neurological phenomena both causing high intracellular calcium: seizures and cortical spreading depolarization (CSD)^7^.

CSD was first documented in 1944, when Aristides Leão reported his first series of electrocorticography recordings of slow propagating waves of depression in rabbit cortex, an investigation he began originally to induce epileptiform discharges^8, 9^. He would later go on to characterize these traveling waves as large scale depolarizing events, exhausting the tissue into a state of depression, hence the term CSD^10^. While CSD is largely considered to be pathologic across a wide array of diseases^11–15^, its relationship with seizures appears to be more complicated ^16^. Observations in animal models^17–20^ and patients^21, 22^ have demonstrated a strong association between the two events with CSD occurring often at the end of seizures. Indeed, the ionic shifts that occur during seizures are conditions that parallel those of CSD. Large extracellular deposition of glutamate and potassium^7, 23^ and the shrinking of extracellular space^24, 25^ create a hyperexcitable state for neurons that can beget CSD, in a feedforward amplificatory fashion^26^. The subsequent impact of CSD in seizures is both deleterious and protective, being implicated as an underlying cause of sudden unexplained death in epilepsy (SUDEP)^27^, as well as a mechanism of seizure termination^17^. Thus, having a better understanding of the interplay between these two neurologic phenomena could be of vital importance for both the suppression of seizures, as well as prevention of one of epilepsy’s most feared complications.

In our previous studies using *in vivo* two-photon calcium imaging in awake mice to evaluate seizure dynamics^28^, we observed slow propagating calcium waves at the end of seizures, which corresponded with an absence of high frequency neuronal firing in electrocorticography (EEG). We hypothesized that these may be CSD, as large calcium transients are a known occurrence during spreading depolarizations^29, 30^ and have been observed as waves^31^ with similar spatiotemporal properties. As the increase in intracellular calcium during CSD exceeds that during seizures^7^, extracellular influx is unlikely to be the only contributing source of calcium, raising the possibility of ER involvement. Furthermore, since elevated calcium levels are hypothesized to mediate seizure termination^32^, this dramatic calcium rise may underly the purported seizure suppressive effects of CSD.

To investigate the mechanisms underlying these stark intracellular increases in calcium, we generated an adeno-associated viral (AAV) vector to transduce neurons with two calcium indicators at the same time, each targeted to a separate intracellular compartment. In the cytosol we express a yellow derivative of the GCaMP family, XCaMP-Y^33^, and in the ER lumen we express a red indicator, RCatchER^34^. RCatchER is a low affinity (*K*_d_ = ∼400 µM) calcium indicator protein with fast kinetics (<1 ms) and a fluorescence intensity that varies linearly with calcium concentration, making it an optimal indicator for ER calcium. We then intravitally recorded generalized seizures^35^ in awake mice, concurrently with EEG and direct current (DC) recordings, with DC being the gold standard for confirmation of CSD.

In the present study we determine that these slow propagating post-ictal calcium waves are in fact CSDs. We next show that CSD is marked by a stark depletion of ER calcium not occurring during normal or epileptiform activity. Our spatiotemporal analysis at a cellular level indicates that this depletion is consistent with a calcium-induced calcium release (CICR). In addition, we observe that depletion of ER calcium also occurs during CSD evoked by electrical stimulation. Finally, we provide causal evidence that naturally occurring CSD is associated with a suppression of post-ictal epileptiform activity and this same suppression can be achieved through the biologically similar electrically evoked CSD. Collectively this work offers new insight into the biological underpinnings of CSD and its potential utility for seizure control.

## Results

### Intravital imaging of cytosol and ER calcium stores in awake mice

To study intracellular calcium dynamics *in vivo* with high spatiotemporal resolution, we developed a recombinant AAV to express two genetically encoded calcium indicators (GECIs) of different colors pan-neuronally through the human synapsin I (hSynI) promoter^36^. The yellow XCaMP-Y^33^ and the red-shifted RCatchER^34^ GECIs were separated by the self-cleaving P2A peptide to obtain similar expression levels (Fig. 1a). While the XCaMP-Y will be localized to the cytosol, the RCatchER includes calreticulin and KDEL sequences, for targeting and retention in the ER lumen, respectively. We confirmed this expression pattern by immunohistochemistry (Fig. 1b). These indicators were selected for their ability to be simultaneously excited with a single wavelength of light between 1000 and 1040 nm (Fig. 1c). Emission was bandpass filtered to isolate each’s distinct signal, thus enabling simultaneous calcium imaging in the two subcellular compartments with single cell resolution. We stereotaxically injected this AAV into the motor cortex of mice and installed chronic cranial windows with head plates to facilitate repeated imaging within subjects. We then performed awake, head-fixed imaging in cortical layers 2/3 (Fig. 1d) to record cytosolic and ER calcium changes across hundreds of neurons in the field (Fig. 1e). During spontaneous locomotion activity, we observed transient increases in cytosolic calcium, classically serving as a proxy for neuronal activity/action potentials, while the ER calcium remained relatively stable (Fig. 1f-i) in contrast to the subsequent recordings during pathologic activity.

**Figure 1.**
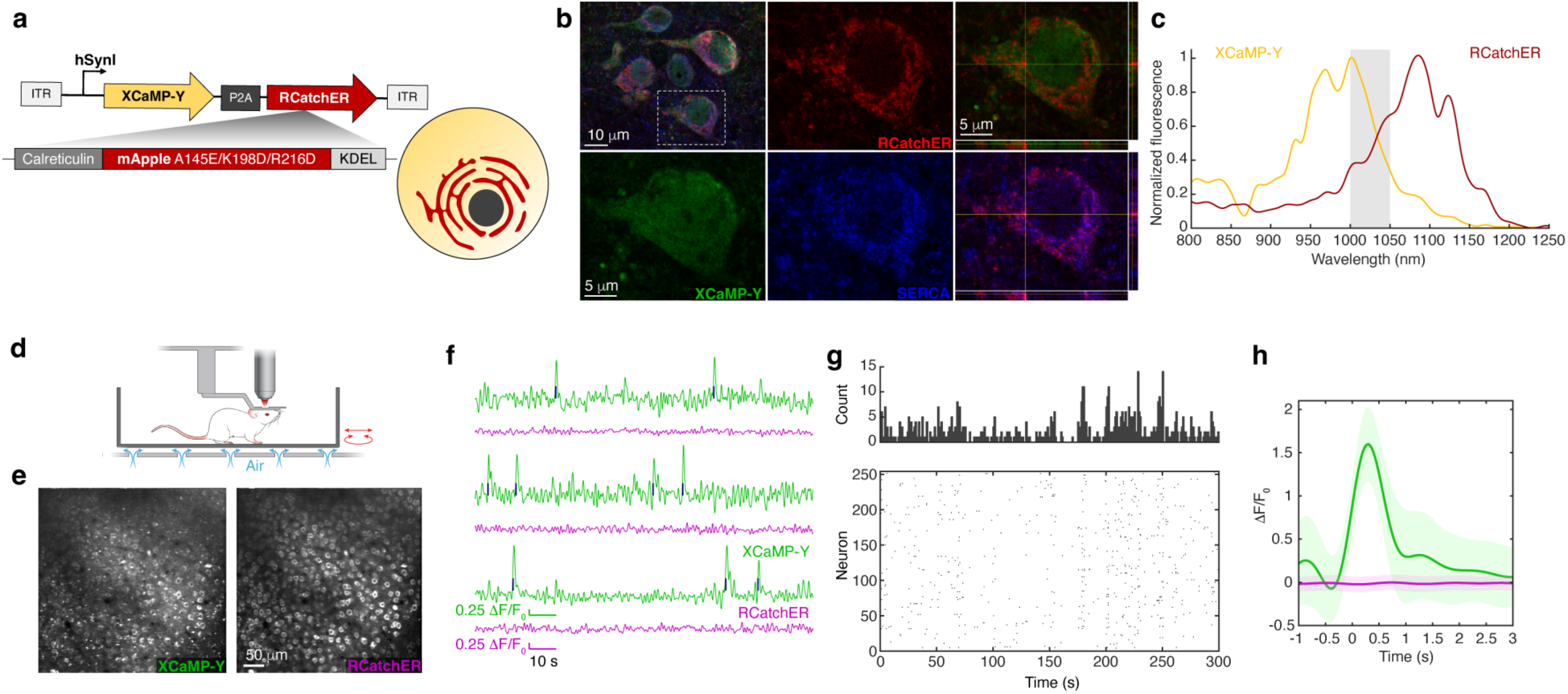
Dual-color two-photon simultaneous intravital calcium imaging of the cytosol and ER during neuronal activity in awake mice. **(a)** Schematic of AAV cassette encoding the XCaMP-Y and RCatchER GECIs (left) enabling mutually exclusive expression in cytosol and ER respectively (right). **(b)** Immunohistochemistry (confocal) demonstrating localization of XCaMP-Y and RCatchER to the cytosol and ER respectively, with RCatchER but not XCaMP-Y colocalized with the ER marker SERCA. Reconstructions of Z axis at crosshairs presented (right). **(c)** Overlapping two-photon excitation spectra of the XCaMP-Y and RCatchER captured *in vivo* (N=1 mouse; n=298 neurons) with the wavelengths used for simultaneous activation highlighted in grey. **(d)** Illustration of awake *in vivo* recording set-up of a head-fixed-mouse poised on air-suspended chamber. **(e)** Representative field of view (cortical layer 2/3; depth: 200 mm from pial surface) showing time-averaged projections of cells expressing both XCaMP-Y (left, green channel) and RCatchER (right, red channel). **(f)** Representative individual cell normalized calcium fluorescence traces (ΔF/F_0_) of XCaMP-Y (green) and RCatchER (magenta), with detected spike times indicated (blue). **(g)** Spike raster and corresponding histogram (1s bin width) of spontaneous firing detected in XCaMP-Y during 5 min of baseline recording (N=1 subject; n=254 neurons). **(h)** Average spike trace relative to indexed spike time, presented with pooled standard deviation of signal (shaded area).

### CSD observed at generalized seizure termination is marked by a unique depletion of ER calcium

For recording the calcium dynamics during seizures and purported CSD, we coupled our dual-color two-photon imaging with simultaneous EEG and DC recording (Fig. 2a). We recorded EEG through chronically implanted electrodes and DC through a glass microelectrode attached to a DC amplifier. To enable this, we fabricated multipaned concentric glass windows with silicone access ports for repeated access to the brain (Fig. 2a and Supplementary Fig. 1). We induced epileptiform activity by subcutaneous administration of pentylenetetrazol (PTZ; s.c. 40-50 mg/kg; N=9 subjects, 30 recordings; Fig. 2a-c). We recorded 23 generalized seizures (Fig. 2b), 4 of which were fatal (analyzed separately in a subsequent section). For the non-fatal seizures, we observed pre-ictal spikes (PIS) across all three recording modalities (Fig. 2d, e), beginning within a few minutes of PTZ injection (234±29 s (mean ± SE); N=19 seizures, 8 subjects). Typically, within 10 minutes (471±53 s), we observed a seizure (length: 21±3 s) on EEG followed by a quiescent post-ictal period. Concurrent calcium transients were observed through the cytosolic calcium indicator, while ER calcium stayed relatively stable. Soon after termination of a seizure, we sometimes observed a large and sustained increase in cytosolic calcium (23±5 s; N=6 seizures, 5 subjects), concomitant with a negative DC shift, a hallmark of CSD (17.03±2.85 mV; N=4 seizures, 4 subjects with DC recording). This calcium change was comparable to that observed during seizures (p=0.688, generalized linear mixed effects model (GLME), N=6 seizures, 5 subjects, 116-371 cells/recording), and greater than that occurring during PIS (p=2.87×10^-21^). We also observed a concurrent large, rapid and sustained depletion of ER calcium which was significantly larger than calcium changes occurring during the seizure or PIS (p=1.88×10^-10^ (CSD to seizure), p=7.01×10^-17^ (CSD to PIS); Fig. 2c-e and Supplementary Fig. 2). PTZ administration did not always induce seizures and CSD (Fig. 2b): it could also induce seizures without CSD, or epileptiform spiking (spike-wave discharges, SWDs) that did not progress to seizure (sub-generalized). We did not observe a comparable change in ER calcium during these events. The large increase in cytosolic calcium and ER depletion was specific to CSD and not a general post-ictal phenomenon, with statistically larger changes in post-ictal calcium during CSD as compared to the same post-ictal period in those seizures without CSD (p= 0.0273 (cytosol), p=9.21×10^-21^ (ER), GLME, N=17 seizures (11 without, 6 with CSD), 8 subjects, 43-371 cells/recording; Fig. 2f). Taken together, we confirmed that the slow propagating cytosolic calcium waves observed at the end of seizures corresponded to CSD with the iconic DC shifts, and a depletion of ER calcium stores.

**Figure 2.**
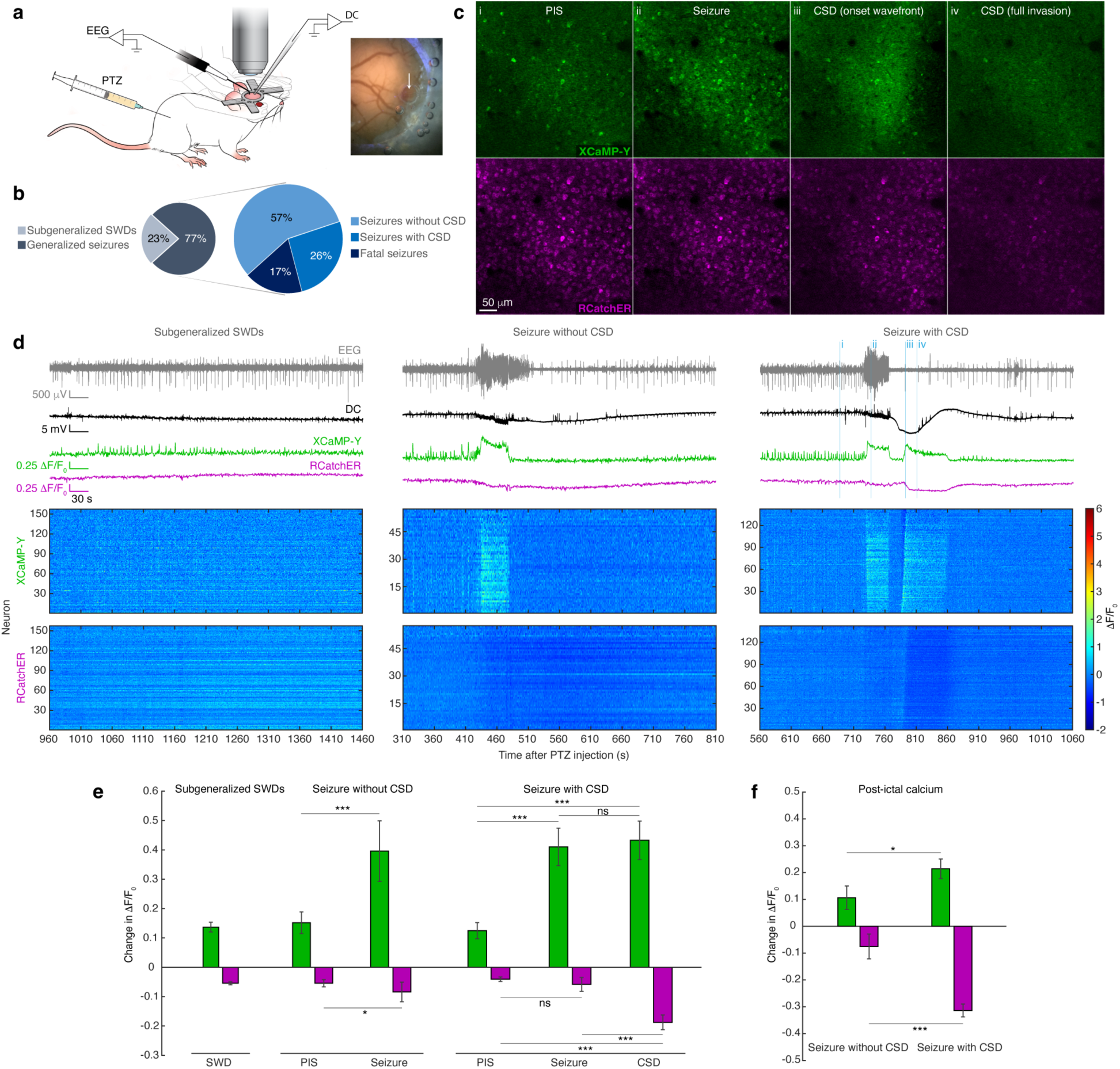
CSD is associated with generalized seizure termination and is marked by a unique depletion of ER calcium stores. **(a)** Illustration depicting the awake head fixed two-photon imaging along with simultaneous EEG and DC recording during PTZ-induced seizures (left). Image of cranial window with glass electrode inserted through silicone access port (white arrow, right). **(b)** Pie charts representing proportion of PTZ injections (N=30) that resulted in generalized seizures (N=23) (left) and, of those generalized seizures, the proportions that occurred with (N=6) or without CSD (N=13) or that were fatal (N=4) (right). **(c)** Representative frames by channel during a PTZ-induced generalized seizure recording depicting the cytosolic (top row) and ER (bottom row) calcium changes during a PIS (i), the seizure (ii) and the CSD (iii: wavefront invasion, iv: CSD after full invasion). Note depletion (left side) of calcium in the recruited area during CSD invasion. **(d)** Mean population calcium fluorescence (XCaMP-Y: green, RCatchER: magenta) with synchronized EEG (grey) and DC (black) recordings during three separate recording sessions within the same subject depicting sub-generalized epileptiform activity and seizures with and without CSD. Corresponding rasters of individual cell calcium transients are presented below (y-axis: neurons ordered from left to right across the field). **(e)** Group level analysis of the average individual cell changes in calcium during each event phase (PIS, seizure and CSD) across each recording type (sub-generalized: N=4 subjects, 7 recordings; seizure without CSD: N=6 subjects, 12 recordings; and seizure with CSD: N=5 subjects, 6 recordings; n=43-413 cells per recording). **(f)** Average post-ictal calcium changes are presented for the seizure recordings with and without CSD. N=17 seizures (11 without, 6 with CSD), 8 subjects, 43-371 cells/recording. The effect of event on calcium levels are modeled using GLME for e and f. Means with pooled standard error are presented in all bar plots. *p<0.05, **p<0.01, ***p<0.001.

### ER calcium depletion during seizure CSD is consistent with a CICR

Having established the sustained post-ictal calcium depletion in the ER to be specific to CSD, we next sought to characterize the spatiotemporal features of these changes. Both cytosolic and ER calcium changes appeared as a wavefront of propagation (Fig. 3a; Supplementary Movie 1). To determine the timing of the changes across the individual cells, we leveraged our slope integral feature detection approach for seizure traveling waves^28^ to find the times individual cells were recruited to the CSD in XCaMP-Y and the time of their ER depletion in RCatchER (Fig. 3b). We generated a colormap depicting relative recruitment times within the channels in the field of view (Fig. 3c). Wavefronts in both channels appeared to move in the same direction, with the wave in the ER slightly delayed.

**Figure 3.**
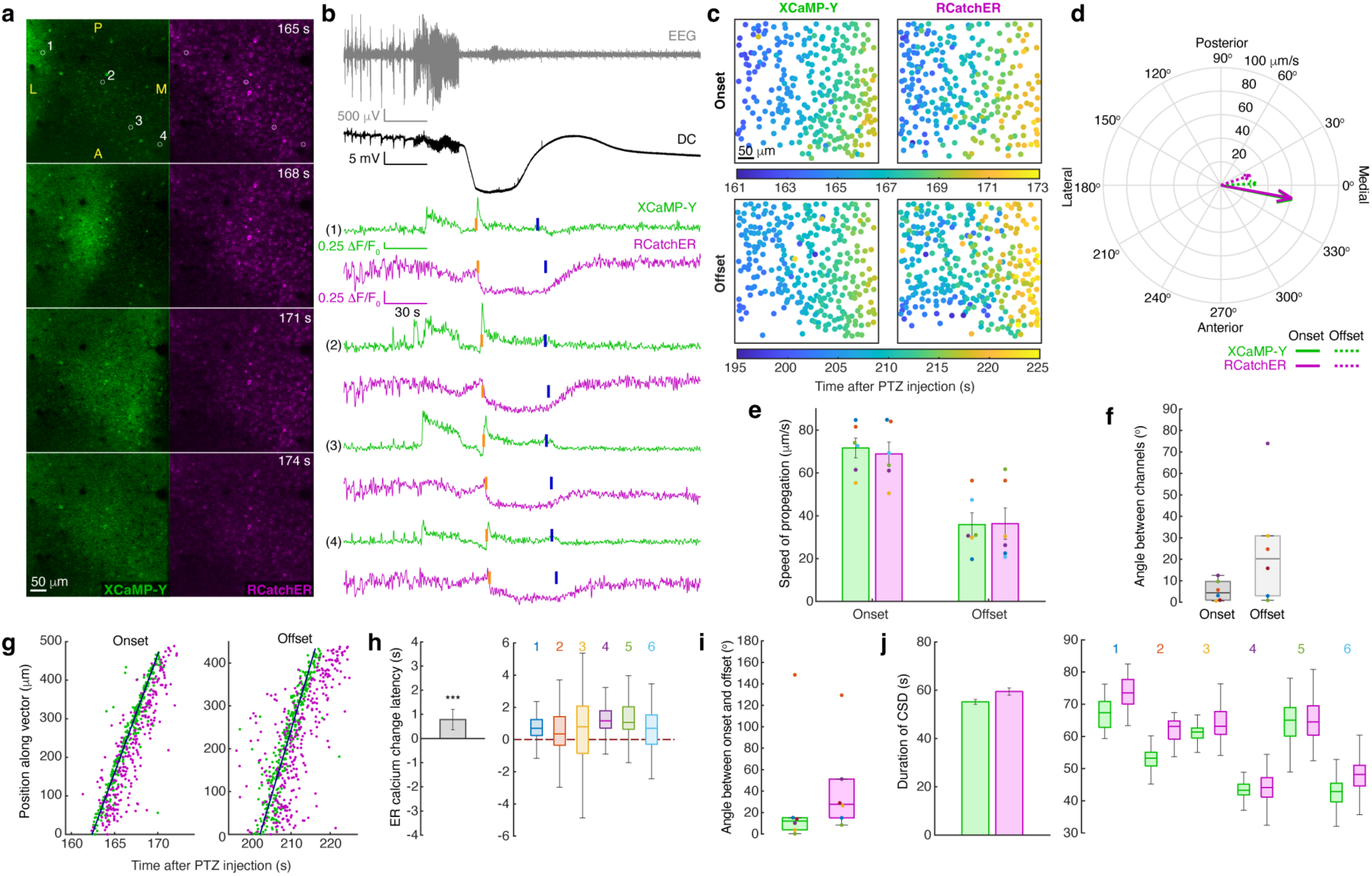
Spatiotemporal dynamics of subcellular compartment calcium changes during CSD. **(a)** Time lapse images depicting the represented CSD wavefront invasion calcium changes. Circles indicate regions of interest (ROIs) of the representative traces in b. Field orientation is indicated (A: anterior, L: lateral, M: medial, P: posterior). **(b)** Representative individual cell calcium traces of XCaMP-Y (green) and RCatchER (magenta), during generalized seizures occurring with post-ictal CSD along with concurrent EEG (grey) and DC (black) traces. The detected recruitment times to the CSD invasion wavefront are indicated in orange and subsequent calcium changes at the offset of the CSD in blue. **(c)** ROI colormap of determined recruitment times for identified cells during the onset and offset of the representative CSD in panel a, with color corresponding to recruitment time. **(d)** Polar plot of the CSD onset and offset propagation vectors modeled by applying spatial linear regression to the neuronal recruitment times shown in panel c, showing wavefront direction (vector angle) and propagation speed (vector magnitude). **(e)** Average speed of CSD propagation at the onset and offset of the event by channel with standard error. Each CSD is depicted using a unique color matched across all the panels in this figure. **(f)** Absolute difference in direction between the XCaMP-Y and RCatchER vectors at the onset and offset of CSD. **(g)** Individual cell recruitment times by channel of the CSD onset and offset projected on to the propagation axis of XCaMP-Y. **(h)** Mean ER calcium change latency relative to the cytosolic calcium change within cell during CSD invasion across all recordings with pooled standard error (left) and the distribution of these latencies for each recording (right). The effect of channel on recruitment times used to compute these latencies are modeled using GLME. **(i)** Absolute difference in direction between the onset and offset vectors of the CSD within channel. **(j)** Average duration of CSD within cell by channel across recordings with pooled standard error (left) and the distribution of the event durations for each recording (right). For group level analysis (e, g, h-j) N=6 seizures across 5 subjects with n=116-371 cells per recording. *p<0.05, **p<0.01, ***p<0.001.

We performed a spatial linear regression^37, 38^ on the recruitment times to determine the wavefront vectors of propagation. We present this first as a polar plot, where the arrows illustrate the direction and speed of the regressed vector of propagation, indicating the overlap in vectors between channels (Fig. 3d and Supplementary Fig. 3). We found that the cytosolic increase and ER depletion propagated in a contiguous fashion, with comparable velocities to each other (Fig. 3d-f). These velocities are consistent with typical CSD velocity^7, 31^. We observed that both cytosolic and ER calcium changes were delayed relative to the CSD DC shift onset (cytosolic: 11.35±0.35 s; ER: 11.77±0.50 s; mean ± SE; N=4 seizures, 4 subjects, 142-371 cells/recording). This is concordant with the DC recording site being about 1 mm lateral to the field of view and thus earlier in the wavefronts’ paths of propagation medially, given the determined velocities of propagation. We next plotted the recruitment times within each channel relative to their projected position along the determined axis of propagation in the cytosolic channel (Fig. 3g). This analysis demonstrated a consistently delayed ER depletion along the same axis of propagation as the cytosolic increase. ER depletion was also found to follow the cytosolic increase within each cell (Fig. 3h), with a significant delay of about 1 second (0.79±0.42 s, p=3.37×10^-8^, GLME, N=6 seizures, 5 subjects, 116-371 cells/recording).

Interestingly, the loss of calcium from the cytosol and return of calcium to the ER also followed a wavefront pattern, occurring in roughly the same direction as the invasion, although with a slower speed (Fig. 3b-g, i; offset). Unlike the initial change in calcium during the CSD invasion, this subsequent change at the end of CSD was more gradual, making the slope integral feature and maximum slope more difficult to discern at the individual cell level. Therefore, we chose to use the point of maximal concavity, the elbow, being the point when a change started. We again found a delay in recruitment relative to the DC shift, consistent with the slower propagation speed of this change (cytosolic: 24.55±0.90 s; ER: 28.15±1.21 s). Computing the time delay between these onsets and offset events within cells, we found lengths of change of calcium comparable between the cytosol and ER (Fig. 3j) and consistent with the average duration of the DC shift (38.93±4.88 s, N=4 seizures, 4 subjects). This short delay in ER calcium release following the same spatiotemporal pattern at the cytosolic increase during CSD, with comparable velocity and duration, suggests a CICR is occurring.

Occasionally the seizure induced was fatal for the subject (Fig. 2b). For these, a terminal spreading depolarization (TSD) was observed following the seizure during isoelectric EEG, with a negative DC shift (-8.84±0.67 s; mean ± SE; N=4 seizures, 4 subjects) that did not recover (Fig. 4a). Additionally, the initial spatiotemporal changes in calcium observed upon TSD invasion were similar to those observed with CSD in the non-fatal seizures. The waves propagated with similar speeds and direction (Fig. 4b-e), with the depletion of ER calcium following the cytosolic increase, albeit with a slightly longer delay (Fig. 4f, g), and smaller magnitude of cytosolic calcium change (Fig. 4h). Notably, the calcium changes did not return to their baseline values, with cytosolic calcium remaining high and ER depleted, consistent with cell death.

**Figure 4.**
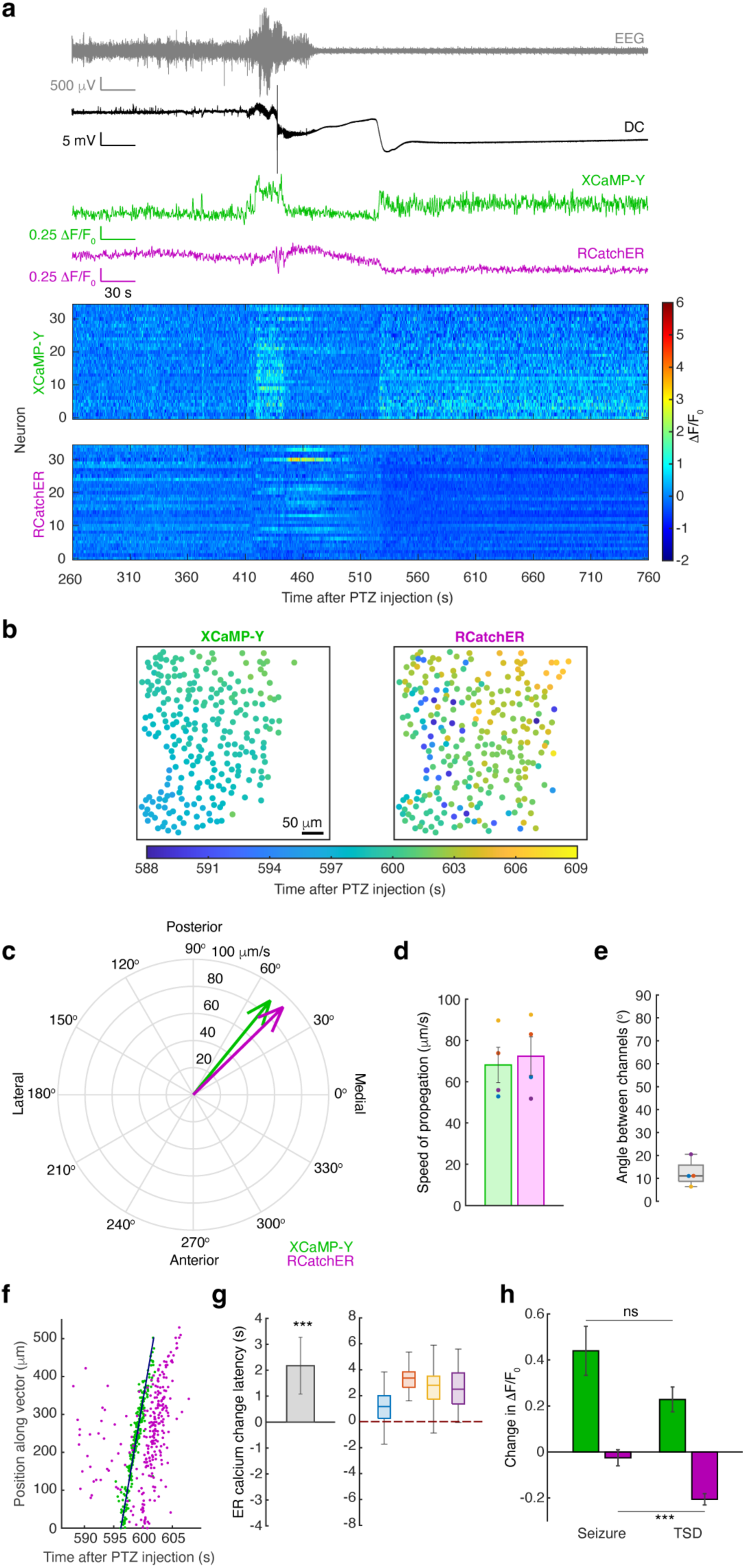
TSD observed during fatal seizures demonstrate a permanent increase in cytosolic calcium and depletion of ER calcium stores. **(a)** Mean population calcium fluorescence (XCaMP-Y: green, RCatchER: magenta) with synchronized EEG (grey) and DC (black) recordings during a fatal generalized seizure occurring with TSD from a representative subject. Corresponding rasters of individual cell calcium transients (y-axis: neurons ordered from left to right across the field) are presented below. **(b)** ROI colormap of determined recruitment times for identified cells during TSD, with color corresponding to recruitment time. **(c)** Polar plot of the TSD propagation vectors by channel, showing wavefront direction (vector angle) and propagation speed (vector magnitude). **(d)** Average speed of TSD propagation by channel with standard error. Each TSD is depicted using a unique color matched with panels e and g in this figure. **(e)** Absolute difference in direction between the XCaMP-Y and RCatchER vectors during TSD. **(f)** Individual cell recruitment times by channel of the TSD projected on to the propagation axis of XCaMP-Y. **(g)** Mean ER calcium change latency relative to the cytosolic calcium change within cell during TSD across all recordings with pooled standard error (left) and the distribution of these latencies for each recording (right). The effect of channel on recruitment times used to compute these latencies are modeled using GLME (p=6.25×10^-11^, N=4 recordings, 4 subjects with N=35-289 cell/recording). **(h)** Average individual cell changes in calcium by compartment during TSD across recordings presented with pooled standard error. The effect of event on calcium level are modeled using GLME (cytosol: p=0.234, ER: p=4.81×10^-4^, N=3 recordings, 3 subjects with N=35-289 cells/recording). For all group level analysis (d, e, g, h) N=7 recordings across 7 subjects with n=82-418 cells per recording. *p<0.05, **p<0.01, **p<0.001.

### ER calcium depletion is a conserved feature across multiple types of CSD

We next sought to determine if these calcium changes were specific to CSD in the context of seizures or if they were a property of CSD itself. For this we used an electrical stimulation model of CSD^39^ where we stimulated through two epidural electrodes chronically implanted adjacent to the recording field, while conducting the same imaging, EEG and DC recording paradigm we employed earlier (Fig. 5a). By applying bipolar stimulation between the two electrodes (2 kHz, 100 µA, square wave, 10 s) we were able to reliably induce CSD (Supplementary Movie 2). The DC shift during electrically evoked CSD was comparable to that during PTZ-induced CSD (Fig. 5b; -19.28±1.97 μV; mean ± SE; *N*=7 recordings, 7 subjects). Similarly, during the electrically induced CSD we observed changes in calcium that paralleled PTZ-induced CSD, both in terms of magnitude (Fig. 5C) and spatiotemporal pattern (Fig. 5d, e), albeit with the CSD propagating at a faster speed, radially from the electrode pair. The cytosolic and ER calcium changes occurred with the same velocities (Fig, 5f, g), with the ER depletion following the cytosolic increase (Fig 5h, i). We observed a delay in the calcium changes relative to the DC shift consistent both with the position of the imaging field relative to the simulating electrodes, and with the speed of propagation (cytosol: 2.74±0.30 s; ER: 3.03±0.41 s). The durations of the calcium changes (Fig. 5j) were again consistent with the average duration of the DC shift (40.53±6.03 s, N=6 recordings, 6 subjects). In addition to the high frequency stimulation of long duration (2 kHz for 10 s), we were able to induce CSD with short trains of stimulation of lower frequency, as typically used in responsive neurostimulation (RNS) devices (Supplementary Fig. 4; N=3 subjects, 2 trials per subject; 250-750uA [charge density within an order of magnitude of RNS range], 200 Hz, biphasic, 160 µs pulse width, five 100-ms trains with 5 s inter-train interval)^40, 41^. Thus, the intracellular calcium dynamics observed during seizure associated CSD were also found with electrically induced CSD, suggesting that the depletion of ER calcium is conserved across CSD.

**Figure 5.**
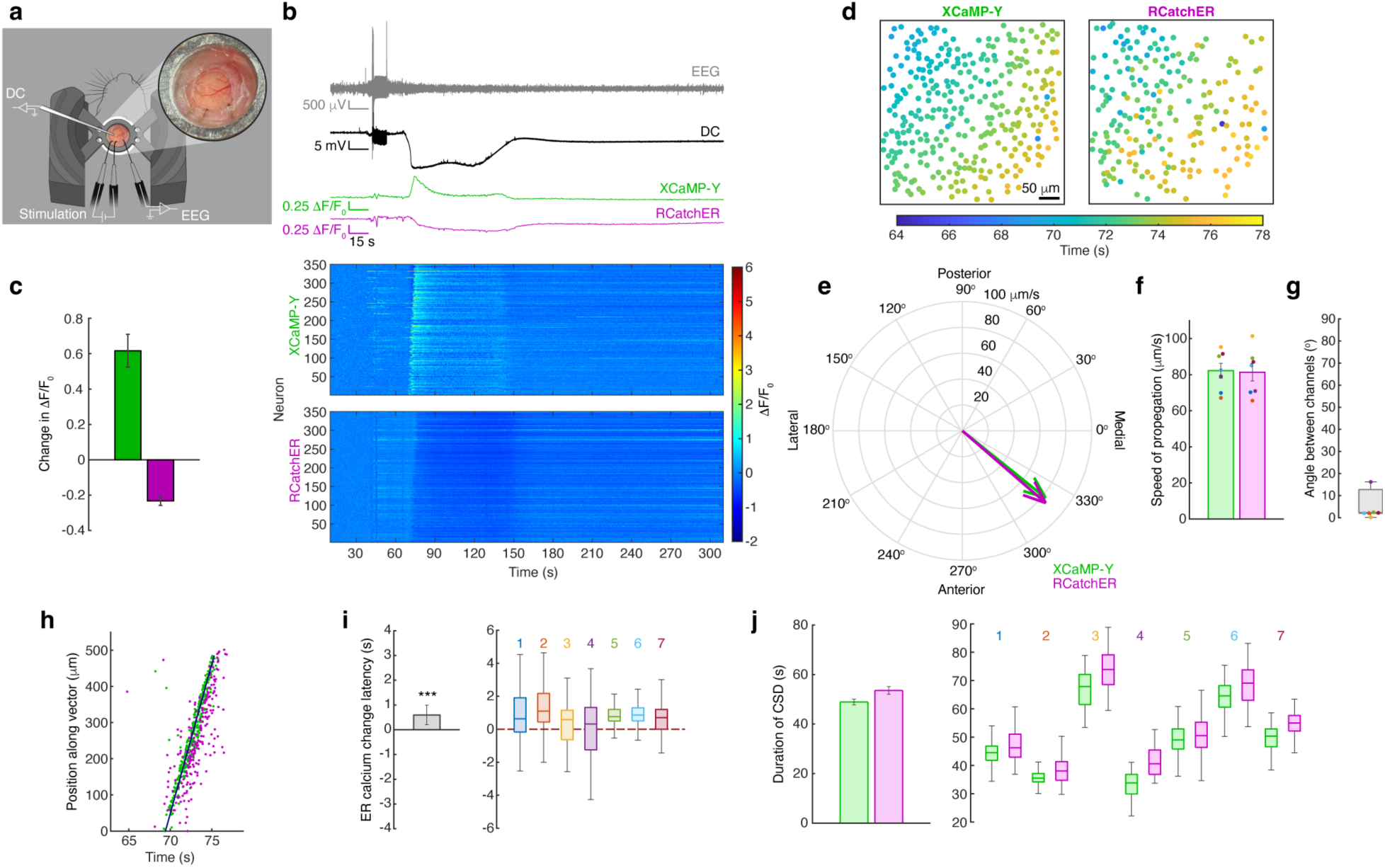
Spatiotemporal subcellular calcium dynamics during stimulation-induced CSD demonstrate ER depletion. **(a)** Illustration depicting the awake head fixed two-photon imaging along with simultaneous EEG and DC recording during stimulation-induced CSD. Expanded is an image of a cranial window with access port and electrodes for stimulation and EEG. **(b)** Mean population calcium fluorescence (XCaMP-Y: green, RCatchER: magenta) with synchronized EEG (grey) and DC (black) recordings during stimulation-induced (10 s) CSD from a representative subject. Note stimulation artifact at 30 s. Corresponding rasters of individual cell calcium transients (y-axis: neurons ordered from left to right across the field) are presented below. **(c)** Average individual cell changes in calcium by compartment during CSD across recordings presented with pooled standard error. **(d)** ROI colormap of determined recruitment times for identified cells during CSD, with color corresponding to recruitment time. **(e)** Polar plot of the CSD propagation vectors by channel, showing wavefront direction (vector angle) and propagation speed (vector magnitude). **(f)** Average speed of CSD propagation by channel with standard error. Each CSD is depicted using a unique color matched across all the panels in this figure. **(g)** Absolute difference in direction between the XCaMP-Y and RCatchER vectors during CSD. **(h)** Individual cell recruitment times by channel of the CSD projected on to the propagation axis of XCaMP-Y. **(i)** Mean ER calcium change latency relative to the cytosolic calcium change within cell during CSD across all recordings with pooled standard error (left) and the distribution of these latencies for each recording (right). The effect of channel on the recruitment times use to compute these latencies are modeled using GLME (p=4.63×10^-8^). **(j)** Average duration of CSD within cell by channel across recordings with pooled standard error (left) and the distribution of the event lengths for each recording (right). For all group level analysis (c, f, g, i, j) N=7 recordings across 7 subjects with n=82-418 cells per recording. *p<0.05, **p<0.01, ***p<0.001.

### Post-ictal activity decreases after CSD

CSD has been previously reported to arrest epileptiform activity in rodents^17^. We next sought to see if that observation held true for our generalized seizures. For this, we quantified spike wave discharges (SWDs) typically observed before and after seizures using our EEG recordings (Fig. 6a). We found that the SWD rate significantly decreased post-ictally when a seizure was followed by CSD, but not when a seizure occurred without a CSD (Fig. 6b; without CSD: p=0.383, N=8 seizures, 5 subjects; with CSD: *p=0.031, N=6 seizures, 5 subjects; Wilcoxon sign-rank test), suggesting a negative correlation between CSD and post-ictal epileptiform activity. To further test if CSD is sufficient to suppress epileptiform activity, we induced CSD using electrical stimulation 5 minutes into the post-ictal phase in a subset of seizures unaccompanied by a CSD (Fig. 6c). We found that, while the post-ictal SWD rate before electrical stimulation was not significantly different from the pre-ictal period (Fig. 6d; Friedman test post-hoc comparison; *p*=0.7593), following electrical stimulation evoked CSD, the SWD rate was significantly decreased (Friedman test with post-hoc comparison: \**p*=0.0356, *n*=4 seizures, 3 subjects). An innate gradual decline in post-ictal SWD rate could not account for this observed difference. We compared the recordings in the same time periods (5-10 minutes post seizure) without electrical stimulation and CSD and found there was no significant difference in SWD rate between the pre-ictal and the post-ictal period (*p*=0.7422; *n*=8 seizures, Wilcoxon sign-rank test). Taken together, CSD, whether it occurs naturally or evoked by electrical stimulation, has potential to diminish post-ictal epileptiform activity.

**Figure 6.**
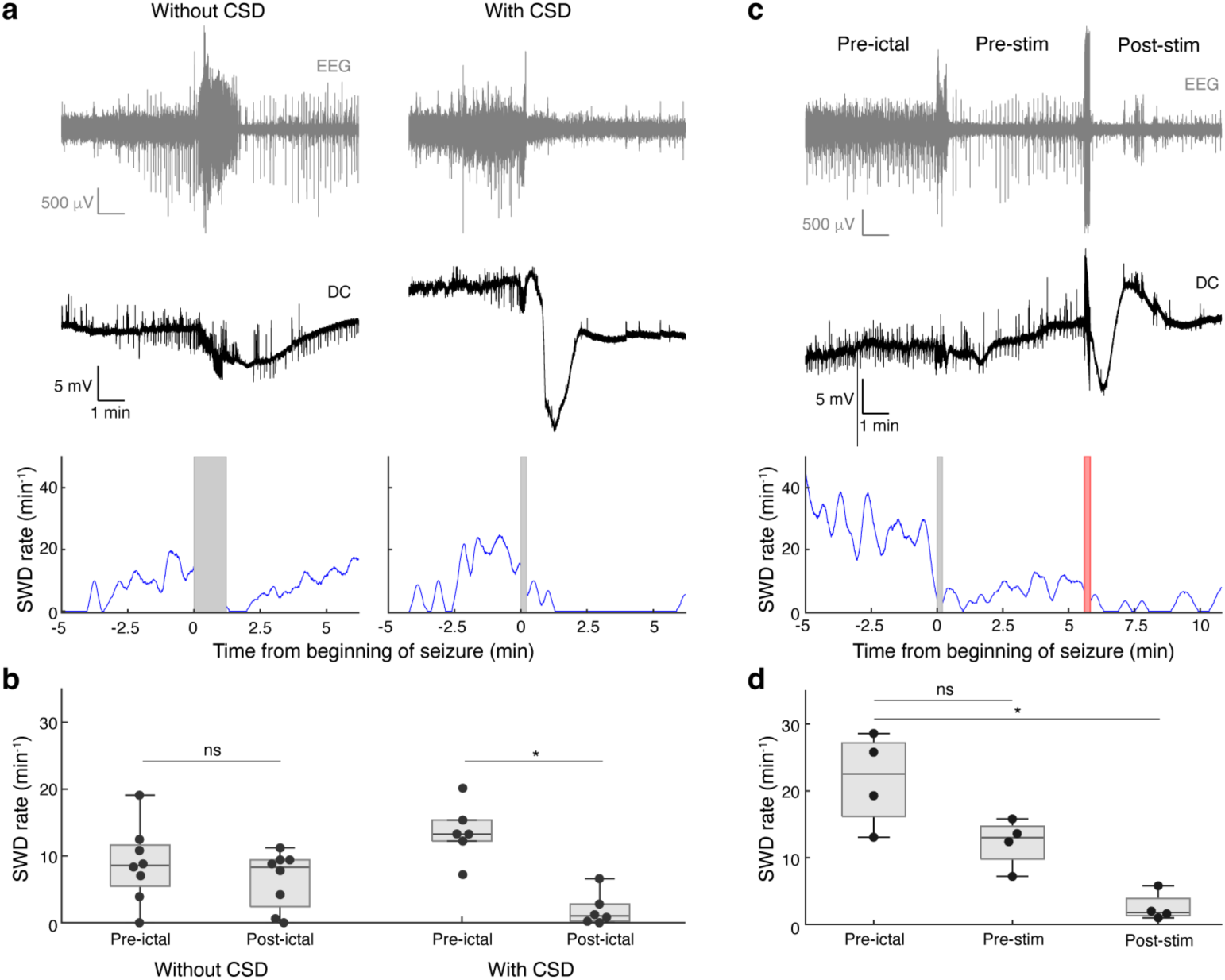
CSD is associated with suppression of epileptiform activity. **(a)** Representative EEG (grey), DC (black) and SWD rates (blue) of the pre- and post-ictal periods for seizures occurring with and without CSD. SWD rate was not calculated during seizure (dark grey box). **(b)** Box plot comparing the SWD rate between the pre- and post-ictal periods with and without CSD. Wilcoxon sign-rank test was used. Without CSD: p=0.313, N=8 seizures, 5 subjects; With CSD: *p=0.031, N=6 seizures, 5 subjects. **(c)** Representative EEG (grey), DC (black) and SWD rate (blue) of the pre-ictal and post-ictal periods, during a seizure without naturally occurring CSD, where a CSD was electrically induced post-ictally. The post-ictal period is further delineated as before (pre-stim) and after stimulation (post-stim). SWD rate was not calculated during seizure (dark grey box) or stimulation (light red box). **(d)** Box plot comparing the SWD rate between the pre- and post-ictal periods (both pre-stim and post-stim) during seizures without naturally occurring CSD, where a CSD was electrically induced. Friedman test with post-hoc comparison was used. \**p*=0.0356, *N*=4 seizures, 3 subjects.

## Discussion

In this study we introduce our XCaMP-Y-P2A-RCatchER imaging construct, for simultaneous *in vivo* two-photon calcium imaging in the cytosol and ER (Fig. 1). RCatchER has previously only been used *in vitro*^34^. Here we expanded the utility of RCatchER by incorporating it into this multi-compartment *in vivo* imaging approach, a novel vertebrate intravital application of calcium imaging to the ER. The ability to capture rapid ER dynamics in awake animals opens vast possibilities for investigators, made all the more accessible through our single AAV design for multicompartment two-color imaging. We envision the usefulness of our construct extending across the biological sciences, from immunology for the study of lymphocyte activation^42^, to cardiology for examining cardiomyocyte contraction^43^.

To demonstrate the utility of *in vivo* RCatchER, we apply this paradigm to a rodent seizure model, which enabled us to uncover ER calcium dynamics unique to CSD. Principally, we observe a depletion of ER calcium occurring during post-ictal CSD (Fig. 2) and electrically induced CSD (Fig. 5) that does not occur during seizures themselves. Depletion of ER calcium was delayed by a few seconds relative to a cytosolic calcium increase, suggestive of CICR (Fig. 3). We observed comparative delays in depletion of ER calcium in TSD (Fig. 4) as well as in electrically evoked CSD (Fig. 5). We also present further evidence of the influence of CSD on seizures, with a focus on the post-ictal suppression of epileptiform activity that correlates with CSD occurrence, which was also recapitulated through electrically evoked CSD (Fig. 6). This suggests post-ictal suppression through CSD may serve as an innate therapeutic mechanism^17^ and also raises the possibility of CSD as a therapeutic electrical stimulation etiology.

Given the potentially fast intracellular dynamics at play during CSD, we selected RCatchER for our paradigm to maximize our temporal resolution, while also permitting simultaneous cytosolic imaging. The vast majority of calcium indicator proteins, including XCaMP-Y, are based on EF-hand motif calcium binding domains, as is the case with the calmodulin of GCaMP^44^, which require cooperative binding and consequently exhibit non-linear fluorescence dynamics^33, 45^. However, RCatchER has a unique calcium sensing mechanism, where a single calcium binding site was engineered on the surface of the scaffold of a red fluorescent protein, mApple^34^. This enables calcium ion binding with 1:1 stoichiometry without cooperativity, whereby its change in fluorescence is not limited by a slow conformational change. Consequently, RCatchER is an exceptionally fast acting calcium sensor, whose dissociation kinetics exceed the temporal resolution of stopped-flow fluorescence measurements, while maintaining fluorescence outputs linear to the calcium levels. This ideally positions RCatchER to capture ER dynamics.

Combining our dual-color imaging approach with simultaneous EEG and DC recording we were able to capture single-cell neural activity and calcium homeostasis dynamics across hundreds of neurons in awake mice during PTZ-induced seizures and subsequent slow propagating calcium waves. It is known that CSD can follow or interrupt seizures^17, 21^. Large calcium increases propagating as traveling waves have also been observed during CSD^31, 46^. Similar calcium increases^47^, and traveling waves^48–50^ have been observed following seizures. While some have inferred that these calcium waves are therefore indicative of CSD, here we offer definitive proof of this association by corroborating these calcium waves with DC recordings.

An association between seizure and CSD occurrence has been demonstrated since early investigations into CSD. Studies from the 1950s in anesthetized rabbit cortex showed that electrically induced after-discharges, as well as PTZ-induced epileptiform activity could be followed by slow potential changes and CSD^51, 52^. The authors hypothesized that the SD was serving to arrest the seizures. They also noticed that the intensity of the epileptiform activity was typically less in tissue that had experienced a prior CSD, indicative of a potential protective consequence of CSD. These ideas are furthered by a PTZ kindling study in rats, where the investigators found that with kindling the occurrence of CSD decreased while the occurrence of epileptiform activity increased^20^. They, too, postulated that the CSD was arresting the seizure, but added that as kindling progressed, between the evolution to a more gradual seizure onset and upregulation of potassium reuptake mechanisms, the increase in extracellular potassium became less abrupt and thus the probability of CSD occurrence decreased, in turn arresting fewer seizures. They also noted a decrease in interictal spiking following CSD, a finding consistent with our results here. Induction of CSD has also been demonstrated to suppress spike wave discharges^53, 54^ and seizures^17^ in animal models, findings concordant with our demonstration of electrically induced CSD decreasing post-ictal spiking (Fig. 6).

The large rise in intracellular calcium we observed during seizures and CSD could itself have implications for seizure termination. During seizures, membrane depolarization opens voltage gated calcium channels and releases magnesium block of calcium-permeable NMDA glutamate receptors (NMDARs), which - coupled with excessive extracellular accumulation of glutamate - causes a large and rapid influx of calcium. This results in acidification of the intracellular compartment through the exchange of calcium and protons across the Ca^2+^/H^+^ ATPase. This acidification, in turn, can lead to decreased conductance of voltage and ligand gated channels^23^. As such, the acidification during a seizure has been hypothesized to promote seizure termination. Additionally, excessive activation of ATPase would contribute to the depletion of ATP, another potential component of seizure termination. Furthermore, the increase in intracellular calcium will lead to excessive neurotransmitter release and eventual depletion of synaptic vesicles, another potential factor in seizure termination^32^. The same processes occur during CSD^55, 56^, although to a much greater extent. The intracellular calcium concentration is estimated to rise to 6-25 μM during CSD^7, 57^, an order of magnitude greater than the rise of calcium during seizures (700 nM)^23, 58^. In this study, we observed a significant release of ER calcium during CSD, but not during seizures (Fig. 2). It is possible that this large ER calcium release specific to CSD is contributing to the higher cytosolic calcium concentration compared to that during a seizure, which could be a factor in the anti-seizure effect of CSD^17^.

Our finding that the depletion of ER stores follows the increase in cytosolic calcium, presumably through voltage gated calcium channels and NMDARs, is suggestive of a CICR occurring within neurons. While RCatchER and XCaMP-Y differ vastly in their kinetics, such a difference in the sensors in and of themselves cannot explain the delay, with the kinetics of RCatchER far exceeding those of XCaMP-Y. Calcium release from ER stores is mediated through two receptor families, ryanodine receptors (RyR) and inositol (1,4,5)-triphosphate (IP3) receptors. While IP3 is implicated in a variety of cell signaling cascades, RyR activation is more specific^3^. The RyR1 isoform, which has minimal expression in nervous tissue, demonstrates voltage dependence, mediated by a mechanical interaction with voltage dependent calcium channels. RyR2, which is the isoform predominantly expressed in the cortex, is activated through calcium influx and is voltage independent^59, 60^. The RyR response to calcium is biphasic, where they are activated at around ∼1 μM of calcium and are subsequently inactivated when calcium exceeds ∼1 mM^61–63^. The calcium dependence of RyR activation may explain our finding that depletion of ER calcium was small and insignificant during seizures, perhaps because intracellular calcium does not reach RyR activation levels, whereas in CSD, it exceeds this threshold. Furthermore, voltage dependent activation occurs faster (∼2 ms) than the CICR, and while the activation of the RyR channel is relatively fast (<10 ms) in the immediate presence of sufficiently high calcium, the widespread induction of CICR is slower, being dependent on the rate of calcium influx and diffusion through the cytosol^2, 63^. IP3 mediated calcium release can also occur over a similar timeframe. As calcium is a co-agonist for the IP3 receptor and considered the driving force behind larger concerted calcium releases, such as the one we observe here, this would also support the CICR hypothesis, albeit by an alternate pathway^64^. Thus, the relative recruitment dynamics we observe during CSD are temporally corroborative with a CICR, although further investigations are needed to identify the precise receptors at play.

Following ER depletion, store operated calcium entry (SOCE) can occur through calcium release activated channels (CRACs) at the cell plasma membrane, including the ORAI1/STIM1 complex, further increasing cytosolic calcium^65–67^. While NMDARs have been considered the primary route of entry contributing to the large increase in cytosolic calcium during CSD^68^, SOCE could be contributing to the sustained increase in cytosolic calcium observed in CSD.

Gain of function mutations causing ‘leaky’ RyR2 have been linked to SUDEP, with knock-in studies of the same mutated receptors in rodents demonstrating decreased threshold for seizures and CSD^69^. While the effects of this mutation could be upstream of CSD by promoting cortical excitability, it could also be directly impacting the generation of CSD, particularly if CICR or SOCE are central to the CSD mechanism rather than only downstream consequences.

While we were able to make a comparison of the timing of cytosolic increase to ER calcium depletion at the start of CSD, it was more difficult to make such a comparison for the offset of the CSD. This is primarily due to the gradual and often smaller calcium changes occurring at the end of the CSD, leading to less precision in the selection of an offset time. However, this variability is small when compared with the length of CSD and with the offset event duration across the population and thus does not dramatically impact the spatial regression for these slow traveling waves. Mechanistically, irrespective of the imprecision in measurement, the timing of these two events are within an acceptable range for the expected timing of cytosolic calcium return to the ER through the sarco(endo)plasmic reticulum ATPase (SERCA) pump^70^.

It is important to note that other ER dynamics are occurring during CSD aside from the observed calcium depletion that could have implications for our findings. ER is known to undergo morphological change during CSD: fission and fragmentation of ER occurs in a calcium calmodulin-dependent kinase II (CaMKII)-dependent manner^46^, which can potentially contribute to the RCatchER signal we measured in this study. However, we deduce that the ER calcium signal we measured in this study was not significantly affected by the morphological change of ER due to the slow nature of ER fission and fusion. The RCatchER signal decreased within a second of the increase in cytosolic calcium (Fig. 3), whereas the ultrastructural changes in the ER were found to occur several seconds after the calcium influx, indicating that the ER calcium release precedes the fission. Additionally, the RCatchER signal recovered within a minute following the depletion of ER calcium (Fig. 3), while the ER fusion should take several minutes to restore continuity of ER. Even if ER is fragmented into beaded structures, that by itself should not hinder the calcium dependence of RCatchER fluorescence. Furthermore, the ER beading occurs predominantly in the neuropil, while our imaging analysis primarily focused on somata. Given the calcium dependence of CaMKII, perhaps the release of ER calcium stores we describe here facilitates the fission event. Another consideration for our finding is the potential impact of intracellular pH on RCatchER’s excitability. While intracellular acidification does occur during seizures, the time course is a gradual change throughout the seizure and continuing during the post-ictal phase^71^, rather than an abrupt drop post-ictally. During CSD acidification also occurs although there is a delay in the drop in pH relative to the DC shift and the pH decrease is sustained for longer than the DC shift^72–74^. Therefore, the dynamics we observe here are not temporally concordant with the pH changes observed. Furthermore, the ER is well buffered, having little change in pH during calcium release and experiences much smaller magnitude pH changes than the cytosol during intracellular acidification^75^.

TSD is an anoxic variant of CSD occurring during death^76, 77^, known to have slightly different mechanisms underlying its calcium dynamics. However, we still observed a depletion of ER calcium stores following a large cytosolic increase, albeit more delayed. Under severe hypoxia, hyperpolarization as a nonspreading depression occurs in the brain, observed as an isoelectric EEG, preserving the ATP stores necessary for recovery. If, however, the hypoxic conditions last more than a few minutes, the ATP stores become depleted, the ion gradients across the membranes break down, and a TSD is, in turn, initiated^7, 55^. While calcium and NMDAR have been found to be necessary for CSD occurrence in normoxic conditions, in the context of hypoxia, CSD can occur in the absence of extracellular calcium and NMDAR function^57, 78, 79^. We observed such TSD calcium waves after about a minute of isoelectric EEG during fatal seizure recordings (Fig. 4). While the propagation patterns were similar to CSDs in surviving animals with respect to overlapping direction and speed, the delay in the ER depletion following the increase in cytosolic calcium was slightly longer during TSD. The increase in cytosolic calcium fluorescence was smaller during TSD than other CSD, perhaps contributing to this longer delay in a CICR. Consistent with cell death and ATP depletion, the cytosolic calcium increase was sustained, and ER calcium was not restored. While cells can typically tolerate the length of depolarization and elevated calcium during a seizure or normoxic CSD^55^, with some neuroprotective effect even having been demonstrated for the intracellular calcium concentrations reached during seizures^80, 81^, the levels experienced during CSD when sustained, as is the case in anoxia and TSD, are generally regarded to be toxic^3, 68^.

Our findings support the plausibility of calcium homeostasis dysregulation during CSD. However, further investigation is needed to determine the necessity of CICR in CSD and its seizure suppressive effect. If CICR proves to be mechanistically involved, evidence of its exact mediators, be it RyR or IP3 dependent, could provide us with new targets for modulation, informing our anti-epileptic arsenal. While CSD may be beneficial in the context of widespread aberrant brain activity during a seizure, it can most certainly have pathologic consequences. Being able to mimic the anti-seizure effect of CSD while avoiding its toxic side effects could offer great therapeutic potential. Alternatively, if CICR rather contributes to the prolonged depolarization and toxic consequences of CSD, prevention of this calcium store depletion during CSD could also be of benefit.

The clinical implications of our investigation afford insight not only to potential neuromodulatory targets, but also to hypothetical mechanisms underlying established therapeutics, namely electrical stimulation used for seizure control. Notably, we were able to induce CSD using different stimulation parameters, including ones paralleling typical RNS settings. Indeed, repeated cortical stimulation has been shown to cause increases in extracellular potassium, likely contributing to CSD induction^82^. The clinical benefits of RNS are likely multifaceted. In large part the decreased seizure incidence is hypothesized to be mediated by neuroplasticity^83^, rather than direct arrest of seizures, as seizures in most patients are not terminated by RNS, and the majority of stimulation occurs interictally. However, in those patients for whom seizures are arrested, these results raise the possibility that CSD could be occurring and contributing to the suppressive effect.

## Materials and Methods

### Molecular Biology

An AAV2 transfer plasmid containing the XCaMP-Y-P2A-RCatchER cassette was generated through standard molecular biology (restriction enzyme [RE] digestion, ligation, transformation, and plasmid purification). The RCatchER cassette in the pcDNA3.1 vector^34^ was cut with NheI and EcoRV and ligated into the NheI and blunted HindIII RE sites in an AAV2 transfer vector with the human synapsin I (hSynI) promoter (Addgene plasmid number: 100843), resulting in pAAV2/hSynI-RCatchER. The coding sequence of XCaMP-Y^33^ (GenBank accession number: MK770163.1) was fully synthesized through a commercial service with an NheI RE recognition site added at the 5’ end (gBlocks gene fragments, Integrated DNA Technologies). The NheI and ClaI fragment of XCaMP-Y was ligated together with the ClaI and BamHI fragment of separately synthetized P2A sequence into the respective RE sites upstream of RCatchER, resulting in pAAV2/hSynI-XCaMP-Y-P2A-RCatchER. The correct sequence was confirmed through RE analysis, as well as Sanger and next-generation sequencing (Eton Biosciences and Plasmidsaurus, respectively). Endotoxin-free DNA was obtained through a commercial midiprep plasmid purification kit (NucleoBond Xtra Midi EF, Takara Bio USA).

### Viral Vector Production

We produced a recombinant adeno-associated viral (rAAV) vector pseudotyped with AAV9 capsid protein in-house following a modified standard procedure^84^. In brief, we transfected human embryonic kidney 293FT cells (Invitrogen) with three plasmids (helper [Addgene plasmid number: 112867], AAV2/9 rep/cap [Addgene plasmid number: 112865], and pAAV2/hSynI-XCaMP-Y-P2A-RCatchER) at 1:1:1 molar ratio using the calcium-phosphate method. We then harvested the rAAV, purified through iodixanol gradient ultracentrifugation, and concentrated in Dulbecco’s phosphate-buffered saline (D8537, Millipore-Sigma) supplemented with 0.001% (v/v) Pluronic F-68 (Millipore-Sigma). We aliquoted and stored it at -80°C until surgery. Using quantitative PCR, we determined the titer to be 4×10^14^ viral genomes/mL.

### Cranial Window Fabrication

To generate windows for chronic imaging with the capability for repeated access via a pulled glass micropipette electrode for DC recording, we adapted protocols for concentric window design^85^ with single plane windows along with silicone access ports^86, 87^. We used two smaller inner layer coverslips and one larger outer layer coverslip (3 mm and 5 mm diameter, respectively, #1 thickness, Warner Instruments), and generated a 0.7 mm diameter access port placed 0.75 mm from the center (Supplementary Fig. 1a). When deciding on diameter of the window it is important to consider the planned angle of penetration for the recording electrode, along with the thickness of the electrode and depth of the access port so as to ensure the window has a large enough diameter to accommodate these constraints (Supplementary Fig. 1b). We began by first etching the holes in each glass layer at the correct coordinates to ensure the holes alignment (0.75 mm from the side of the 3mm glass and 1.75 mm from the edge of the 5mm glass) when the glass layers are stacked concentrically. For etching we suspended the glass using a spring hinged clip to apply light, but sufficient pressure to the edges of the cover glass. To protect the glass from damage, we covered the clip with heat shrink wrap. Using a conical sharp tip fine grit stone grinding burr (CA1063, Minimo Precision Instruments and Tools) with a dental handpiece at a medium speed (∼6000-7000 rpm), we slowly hand dry etched each cover glass at a 45° angle, moving halfway through each piece of glass from each side to meet in the center, with applying dust free air routinely to clear away the silicone dust during etching. Upon meeting in the center, we moved the burr into a vertical position (perpendicular to the cover glass) to round out and straighten the beveled edges from both sides of the glass as needed to minimize imaging artifacts. We then cleaned each cover glass with lens paper and compressed air.

We next assembled the cover glass using optical adhesive (Optical Adhesive 71, Norland), on a Styrofoam platform with vertical pins (000 insect pins; 0.5 mm shaft diameter) to thread the holes for cover glass alignment. We adhered the layers serially using a small amount of adhesive (just enough to spread to the edge of the 3mm glass), first the 5mm to a 3mm and then the second 3mm to the first, with interleaved UV curing (365 nm; wavelength of peak absorbance) for 30 min. We then cured overnight (>12 h) at 50°C. The optical adhesive must be uniformly distributed between articulating surfaces to prevent thin film interference.

The final step is to prepare the silicone membrane by filling the access port with optically transparent silicone (Sylgard 184, Dow Corning), selected because it can be cured at room temperature given the initial limited temperature range tolerance of the optical adhesive (-15-60°C). We prepared the silicone at a 10:1 mixture by weight of base to catalyst. Next, we increased the viscosity of the silicone through heating to facilitate easier application to the port using heat gun at 150-200°C for about a minute. Then using a 30-gauge insulin needle we applied a tiny drop of the prepared silicone to fill the access port of the windows while suspending them in mid-air by their edges with light tension to prevent wicking. We then transferred the suspended windows to a vacuum chamber and cured at room temperature for two days to prevent bubble formation in the silicone. We then followed this with a one-day cure at 50°C. Curing at a higher temperature increases the silicone’s strength (higher shear modulus), while curing at a lower temperature increases its elasticity (lower strain at break)^88^. We designed this approach to ensure a balance of strength and elasticity such that the membrane will not deform under increased intracranial pressure (strength), while at the same time will properly re-seal upon needle withdrawal (elasticity). We stored these windows at room temperature. For a full cure we waited an additional four days and implanted them within a few months, before the silicone dried out, becoming more brittle and losing its elasticity.

### Stereotaxic Surgery

All procedures involving live animals were conducted with approval from and in accordance with Emory University’s Institutional Animal Care and Use Committee. Adapting standard protocols for concentric cranial window implantation^85, 89^, we performed stereotaxic cranial window surgery on adult (≥ 90 day-old) albino C57BL6/N male mice (B6N-*Tyr^c-Brd^*/BrdCrCrl, Charles River, Strain Code 493) concurrently with intracortical delivery of rAAV, electrode placement for recording and stimulation, and headplates affixation. In brief, we secured the mice in a stereotaxic frame and maintained them under anesthesia (1.5% isoflurane balanced in oxygen (1 L/min)). We then performed a 3 mm craniotomy over the primary motor cortex. Two injections of rAAV (500 nL each; 2 nL/s) were performed through a pulled glass capillary tube (Nanoject 3.0, Drummond) at 300 μm and 600 μm deep to the pial surface (0.30 mm anterior and 1.75 mm lateral to Bregma)^90, 91^. We then epidurally placed a thin polyamide insulated tungsten wire electrode with exposed tip (125 μm; P1Technologies) at the posteromedial edge of the craniotomy, along with ipsilateral reference and contralateral ground stainless steel screw electrodes (E363/96/1.6/SPC, P1Technologies) in the skull over the cerebellum (0.7 mm burr holes). If the mouse was to be used for stimulation as well, we also placed two additional wire electrodes (same material and size) epidurally about half a millimeter apart along the posterolateral edge of the craniotomy. We prefabricated all electrodes with gold pins to facilitate easy attachment to a recording preamplifier or stimulus isolator. We next placed a cranial window to plug the craniotomy and affixed the window and electrodes to the skull using dental acrylic (C&B Metabond, Parkell) along with a stainless steel headplate (Models 3 and 4; Neurotar). We closed the skin to headplate using tissue adhesive (Vetbond, 3M).

### Two-Photon Imaging and Electrophysiology

We performed resonant scanning (30 Hz, 512x512 pixels) two-photon imaging on mice during acute induced seizures. We used a two-photon microscope (HyperScope, Scientifica) equipped with a pulsed tunable infrared laser system (InSight X3, Spectra-Physics) and controller software (ScanImage, Vidrio Technologies). We selected 1000-1010 nm wavelengths for excitation (Fig. 1C) and separated the emissions by a dichroic mirror (565LP, Chroma) with band pass filters (ET525/50m-2p and ET620/60m-2p, Chroma), collecting the light using GaAsP and multi-alkali red-shifted photomultiplier tube, respectively. Beginning one month following surgery to allow for adequate GECI expression, we head-fixed the mice in a carbon fiber airlifted chamber (Mobile HomeCage, Neurotar), positioned under a long working distance 16x objective (water-immersion, N.A. 0.80, Nikon). We connected the EEG electrodes to an AC preamplifier and a data acquisition system (Sirenia, Pinnacle Technologies). For DC recording, on the day of a recording session, a reference electrode (1-mm diameter Ag/AgCl pellet, Model E205, Harvard Apparatus) was placed in the nuchal muscle of the mouse under isoflurane anesthesia. A micromanipulator (Kopf Instruments) was affixed to the crossbar of the headplate holder. We stereotaxically inserted a long-shank pulled glass electrode (1 mm diameter with ∼70 μm at the tip, 1 – 3 MΩ when filled with the following solution: 125 mM NaCl, 2.5 mM KCl, 1.25 mM NaH_2_PO_4_, 26 mM NaHCO_3_, 20 mM d(+)-glucose, 2 mM CaCl_2_, and 1.3 mM MgCl_2_) through the access port at a 45° angle into the cortex to a depth of 100-300 μm below the pial surface. DC signals were recorded through a patch clamp amplifier, a digital data acquisition system, and software (MultiClamp 700B, Digidata 1550B, and pClamp, respectively, Molecular Devices). Both EEG and DC signals were recorded at 2 kHz, with a 0.5-300 Hz bandpass filter for the EEG and with a 500 Hz low-pass filter for the DC recordings. EEG was continuously recorded while DC recording and two-photon imaging were triggered by a TTL pulse generated in the EEG recording system immediately before either PTZ injection or electrical stimulation.

### Acute Seizure Model

We subcutaneously injected PTZ (40-50 mg/kg; P6500, Millipore-Sigma; sterile saline diluent) and recorded for 20-45 min depending on the course of the seizure. All but one PTZ injection resulted in at most one generalized seizure per recording session. We performed multiple seizure experiments within the same subject provided the seizures were not fatal, with sessions separated by at least a week to circumvent the effects of kindling^92^.

### Electrical Stimulation

We performed electrical stimulation using a waveform and function generator (EDU33210A Keysight, USA) to drive a stimulus isolator (DS4 or DS5, Digitimer, UK) attached to the chronically implanted electrodes. Typical stimulation parameters were 50% duty cycle bipolar pulses with amplitudes of ±200 µA at 2 kHz for 10 s. We stimulated the mice 30-40 s following the start of image acquisition and would acquire 10-15 min of data depending on the course of the CSD. To evaluate the threshold for inducing CSD with RNS style parameters we created waveforms in MATLAB and then used the generator to drive them. We began stimulation at the lowest setting and proceeded to increase current with each subsequent stimulation periods (25, 50, 100, 250, 500 and 750 µA) until CSD was induced, interwoven by 2-minute washout periods. Following CSD we had a 20 min washout before the trial was repeated within the same subject.

For post-ictal simulation-induced CSD experiments we performed electrical stimulation using the same approach as above in a subset of animals during seizures occurring without CSD. In these recordings we waited 5 min following the end of the seizure to ensure a CSD did not occur naturally and to acquire a baseline post-ictal spiking. We then stimulated to induced CSD and recorded for at least an additional 5 min.

### Histology

Following experiments, all subjects underwent transcardial perfusion (4% paraformaldehyde (PFA) in phosphate buffered saline (PBS), 4°C; 32% PFA stock solution, Electron Microscopy Solutions, cat no. 15714) and their brain tissue was extracted. The brains were further fixed overnight in 4% PFA (4°C) and then cryoprotected for 36 hours in 30% sucrose (in PBS, 4°C). The tissue was serially sectioned using a freezing microtome (Spencer Lens Co. equipped with a Physitemp BFS-40MPA Controller and platform) at 40 μm thickness (coronal) and stored in PBS at 4°C.

For triple immunofluorescence, free-floating sections were rinsed in PBS, blocked in PBS solution containing 4% normal donkey serum (NDS), 4% normal goat serum (NGS) and 0.1% Triton-X for 30 minutes at room temperature. After rinses in PBS, sections were then incubated overnight at 4°C with a combination of chicken anti-GFP (1:100, GFP-1020, Aves Labs, Davis, CA, USA), mouse-anti-mCherry (1:200, AE002, Abclonal, Woburn, MA, USA), and rabbit anti-SERCA2 (1:50, A1097, Abclonal, Woburn, MA, USA) in PBS containing 2% NDS and 2% NGS. Sections were rinsed in PBS and incubated in a PBS solution containing Alexa Fluor 488-conjugated goat anti-Chicken IgG (1:1000, Invitrogen A-11039, ThermoFisher Scientific, Waltham, MA, USA), Alexa Fluor 594-conjugated donkey anti-mouse IgG (1:1000, 715-585-150, Jackson Immunoresearch Laboratories, West grove, PA, USA) and Alexa Fluor 647-conjugated goat anti-rabbit IgG (1:500, 111-607-008, Jackson Immunoresearch Laboratories, West grove, PA, USA) secondary antibodies and 2% NDS/2% NGS for 1 hour at room temperature. Sections were rinsed with PBS, then mounted on glass slides, dried and cover slipped with hard set mounting medium containing the nuclear marker, DAPI (Vectashield, H-1500, Vector Laboratories, Burlingame, CA, USA). Images were captured using a Leica SP8 upright confocal microscope and LASX software. Immunofluorescence image processing, including projections and orthogonal view generation, was performed using FIJI^93^.

### Image Processing

We performed imaging pre-processing using the Suite2P software package^94^ with integrated Cellpose^95^, performing motion registration, region of interest (ROI) detection and calcium transient extraction of soma and surrounding neuropil. We additionally manually curated the candidate ROIs to ensure they were from cell bodies and not overlapping with any major blood vessels. We background subtracted the raw fluorescence traces taking the global minima of the neuropil transients for each recording as a proxy for the background signal. We also generated clean somatic signal by subtracting 70% of the surrounding neuropil signal from the somatic ROI signal to remove out-of-plane contamination^94^ and adjust for photobleaching. We normalized the traces as ΔF/F_0_, using the mean of the first 30 s of recording as baseline fluorescence (F_0_). Finally, we double-reverse filtered the traces using an adaptive filtering routine (1 Hz, Butterworth lowpass filter, order 3 to 5) that preserves the phase of the signal prior to processing.

### Data Analysis

For baseline recording spike detection we employed a peak detection heuristic to select all transients at least four standard deviations above baseline. We computed the average spike trace with pooled standard deviation, aligning all the detected transients by their point of maximum slope. The change in calcium level during each spike was taken as the difference between the average signal during half-second windows before and after the recruitment time.

For recruitment time detection during seizures, we used our previously reported automated approach^28^. In brief, the approach uses the mean population calcium trace along with EEG (in this study we also used DC) to find PIS, seizure and CSD onsets in the recordings and defines detection windows around these events. It then searches the individual traces within these windows for local maxima in the integral of their slope, indicative of rapid and sustained increases in cytosolic calcium that occur during recruitment to these seizure related events. These increases must exceed an event and channel specific threshold above baseline for the cell to be considered recruited to the event. All individual cell recruitments are then indexed by the point of steepest slope, being generally accepted as a recruitment time for seizures^96, 97^, and the point least likely to be perturbed by filtering. For processing the RCatchER signal, we inverted the transients, enabling our algorithm, originally designed to find increases in cytosolic calcium, to find the decreases in signal. With the majority of recordings and events we used the clean somatic traces for detection. We also adapted this method to determine recruitment times of CSD offset. However, due to lower signal-to-noise ratio during the offset of CSD and during TSD, we needed to use the somatic traces before neuropil subtraction for event detection. Additionally, for determining the offset of CSD a few modifications to the approach were implemented given the gradual rather than rapid nature of the change being measured. Namely we filtered the traces to 0.05-0.1 Hz and searched for the local minima/maxima of concavity to estimate the beginning of the offset calcium change.

To determine the magnitude of calcium changes in individual cells during seizures and CSD (including TSD), we computed the difference between the average signal during ten and five second windows, respectively, before and after the recruitment times. For the calcium change during the recruitment to PIS, we also computed the difference in calcium levels before and during the spike. However, we found that using the average signal diminished the amplitude of change too much and using only the maxima left the values too subject to shot noise. Therefore, we opted to further filter the traces to 0.5 Hz before taking the maxima during a one second window before the spike to compare with a two second window encompassing the full spike. To determine the impact of event (PIS, seizure or CSD) on calcium change we model the calcium changes using a GLME with effects coding and with each recording session modeled as a random effect. Then to evaluate the if there was a sustained post-ictal calcium change in an unbiased way, we determined the average post ictal calcium signal in each cell over a sliding 15 s window for each seizure recording. Then using the same GLME approach we modeled the impact of CSD on the post-ictal calcium.

For determining vectors of propagation, as we previously demonstrated^28^ we applied a spatial linear regression algorithm, with L1 regularization (originally developed to model interictal events in human intracranial data)^37, 38^, to our data, using the positions of the cells withing the field and the determined event recruitment times. We only considered statistically significant vectors (p<0.05; although for nearly all of the vectors p<0.001) in our analysis, where p-values were computed by comparing model residuals to spatially shuffled data sets, as in prior work ^37^. Speed and direction of propagation were taken from these vectors for comparisons within and across channels. The same recruitment times used to determine these vectors were also used to compare the relative timings of recruitment to these events across channels, modeling the effect of channel on recruitment time using a GLME with effects coding and with each recording session modeled as a random effect.

For computing the DC shift onset and offset we first smoothed the DC trace using a rolling average, downsampling and interpolation. We took the local minima near the DC fall and local maxima near the DC rise of the second derivative of the trace as the onset and offset times, respectively. We used these times to compute the DC shift period lengths and latencies with respect to the calcium changes.

We produced all movies and time-lapse representative frames from the motion registered image stacks output of Suite2P processed using FIJI^93^. We filtered the stacks using a 3D gaussian filter (X-Y: 0.5 standard deviation (SD); time: 1 SD) and down sampled to 6 Hz (movie) or 1 Hz (representative frames) using bilinear interpolation.

To examine the SWD rates in EEG, we used a bandpower-based threshold detection method^28, 98^ to find all EEG spikes in a recording. We then divided the recordings into the specific periods we wished to compare. We used the same pre-ictal period as in the calcium data. We then used Welch’s power spectral density to compute a spectrogram and used specific frequency features to define the boundaries of the post ictal time periods being compared. The post-ictal period began at the end of the seizure, defined as the point when the total power (<100 Hz) fell below 5% of the maximum power achieved during the seizure. For recordings without post-ictal stimulation, we ended the period 5 min later, being equivalent to when we would induce CSD in the post ictal stimulation recordings. For recordings with post-ictal stimulation, we defined the end of the post-ictal/pre-stimulation period as the point where a stimulation artifact EEG power (50-70 Hz & 170-190 Hz) crossed 50% of its maximum power. The post-ictal/post-stimulation period began when the stimulation artifact power fell below 50% of its maximum and ended 5 min later. We computed SWD rates by dividing the totals spike counts during each of these periods by their length of time. We then computed moving-average SWD rate curves (Fig. 6 a-b) by convolving a 30 s-wide Gaussian window with a binarized array of detected spike times at the original signal sampling rate. Statistics were performed using non-parametric pairwise tests, namely the Wilcoxon sign-rank test for comparisons of two groups and the Friedman test for comparisons of more than two groups.

### Two-photon Excitation Spectra

For determining the excitation spectra of the two indicators, we imaged a mouse prepared for *in vivo* imaging, expressing both XCaMP-Y and RCatchER, using the same approach as above, except with galvo scanning (1.07 Hz, 512x512 pixels). We collected images across a series of wavelengths (800-1250 nm; 10 nm interval) in both channels simultaneously (10 frames per wavelength; saved as a time averaged projection). The laser attenuation was calibrated to maintain constant power at the sample across the spectra, adjusting for wavelength-dependent laser output and attenuation variance. Laser power and PMT gains were calibrated to prevent saturation of PMTs at peak excitation values (XCaMP-Y: 970nm, RCatchER: 1100 nm), while ensuring sufficient signal could be observed at 1010 nm. We recorded pre- and post-recording power measurements and calibration frames to verify that power did not attenuate over the course of the experiment, photobleaching did not occur and there was no degradation of the photodiodes. For processing, we concatenated the image series using FIJI and performed ROI detection and transient extraction using Suite2P. We background subtracted the traces (including removing autofluorescence contamination, likely due to the older age of the mouse used), normalized these to maximum power and performed a cubic interpolation between the discrete emission values to produce the spectra curves.

## Acknowledgments

We thank Thomas Eggers for his assistance with generating stimulation waveforms, Henry Skelton for his assistance with confocal imaging, Alexandra Nazzari for her comments on the abstract and Bona Kim for her illustration work. This work was supported by funding from the NIH [F31NS115479 (MAS), R21NS112948 (REG), S10OD021773 (KB)] and the Mirowski Family Foundation (REG).

## Author contributions

Conceptualization: MAS, KB, REG; Methodology: MAS, KB; Software: MAS, ERC; Validation: MAS, KB, ERC; Formal analysis: MAS, ERC, KB; Investigation: MAS, KB, CAG; Resources: REG, KB, JJY, CAG, MAS; Data curation: MAS, ERC, KB; Writing—original draft: MAS; Writing—review and editing: MAS, KB, ERC, CAG, JJY, REG; Visualization: MAS, KB, ERC, CAG; Supervision: REG, KB; Project administration: MAS; Funding acquisition: REG, KB, MAS

## Data and materials availability

The viral vector plasmids generated from this project will be made available to researchers upon request through a material transfer agreement. All data needed to evaluate the conclusions in the paper are present in the paper and/or the Supplementary Material. Code and summary data needed to replicate figures in the paper are publicly archived at Zenodo [repository pending, will be updated prior to publication]. Updated versions of the code will be available at the GitHub repository: https://github.com/Stern-MA/RCatchER_CSD.

## References

1. Mekahli, D., Bultynck, G., Parys, J., De Smedt, H. & Missiaen, L. Endoplasmic-Reticulum Calcium Depletion and Disease. Cold Spring Harbor Perspectives in Biology 3 (2011).

2. Verkhratsky, A. & Shmigol, A. Calcium-induced calcium release in neurones. Cell Calcium 19, 1–14 (1996).

3. Ghosh, A. & Greenberg, M.E. Calcium signaling in neurons - molecular mechanisms and cellular consequences. Science 268, 239–247 (1995).

4. Berridge, M. Neuronal calcium signaling. Neuron 21, 13–26 (1998).

5. Reddish, F., Miller, C., Gorkhali, R. & Yang, J. Calcium Dynamics Mediated by the Endoplasmic/Sarcoplasmic Reticulum and Related Diseases. International Journal of Molecular Sciences 18 (2017).

6. Suzuki, J., Kanemaru, K. & Iino, M. Genetically Encoded Fluorescent Indicators for Organellar Calcium Imaging. Biophysical Journal 111, 1119–1131 (2016).

7. Dreier, J.P. & Reiffurth, C. The Stroke-Migraine Depolarization Continuum. Neuron 86, 902–922 (2015).

8. Leao, A.A.P. Spreading depression of activity in the cerebral cortex. Journal of Neurophysiology 7, 359–390 (1944).

9. Somjen, G.G. Aristides Leao’s discovery of cortical spreading depression. Journal of Neurophysiology 94, 2–4 (2005).

10. Leao, A.A.P. Further Observations on the Spreading Depression of Activity in the Cerebral Cortex. Journal of Neurophysiology 10, 409–414 (1947).

11. Hills, K.E., Kostarelos, K. & Wykes, R.C. Converging Mechanisms of Epileptogenesis and Their Insight in Glioblastoma. Frontiers in Molecular Neuroscience 15, 23 (2022).

12. Dohmen, C., et al. Spreading depolarizations occur in human ischemic stroke with high incidence. Annals of Neurology 63, 720–728 (2008).

13. Lauritzen, M., et al. Clinical relevance of cortical spreading depression in neurological disorders: migraine, malignant stroke, subarachnoid and intracranial hemorrhage, and traumatic brain injury. Journal of Cerebral Blood Flow and Metabolism 31, 17–35 (2011).

14. Strong, A.J., et al. Spreading and synchronous depressions of cortical activity in acutely injured human brain. Stroke 33, 2738–2743 (2002).

15. Curry, R.N., et al. Glioma epileptiform activity and progression are driven by IGSF3-mediated potassium dysregulation. Neuron 111, 682-+ (2023).

16. Kramer, D.R., Fujii, T., Ohiorhenuan, I. & Liu, C.Y. Interplay between Cortical Spreading Depolarization and Seizures. Stereotactic and Functional Neurosurgery 95, 1–5 (2017).

17. Tamim, I., et al. Spreading depression as an innate antiseizure mechanism. Nature Communications 12, 1–15 (2021).

18. Calia, A.B., et al. Full-bandwidth electrophysiology of seizures and epileptiform activity enabled by flexible graphene microtransistor depth neural probes. Nature Nanotechnology 17, 301–309 (2022).

19. Aiba, I., Ning, Y. & Noebels, J.L. A hyperthermic seizure unleashes a surge of spreading depolarizations in Scn1a-deficient mice. Jci Insight 8, 19 (2023).

20. Koroleva, V.I., Vinogradova, L.V. & Bures, J. Reduced incidence of cortical spreading depression in the course of pentylenetetrazol kindling in rats. Brain Research 608, 107–114 (1993).

21. Dreier, J.P., et al. Spreading convulsions, spreading depolarization and epileptogenesis in human cerebral cortex. Brain 135, 259–275 (2012).

22. Fabricius, M., et al. Association of seizures with cortical spreading depression and peri-infarct depolarisations in the acutely injured human brain. Clinical Neurophysiology 119, 1973–1984 (2008).

23. Raimondo, J.V., Burman, R.J., Katz, A.A. & Akerman, C.J. Ion dynamics during seizures. Frontiers in Cellular Neuroscience 9, 14 (2015).

24. Traynelis, S.F. & Dingledine, R. Role of Extracellular-Space in Hyperosmotic Suppression of Potassium-Induced Electrographic Seizures. Journal of Neurophysiology 61, 927–938 (1989).

25. McBain, C.J., Traynelis, S.F. & Dingledine, R. Regional Variation of Extracellular-Space in the Hippocampus. Science 249, 674–677 (1990).

26. Somjen, G.G. Ions in the Brain: Normal Function, Seizures, and Stroke (Oxford University Press, New York, 2004).

27. Aiba, I. & Noebels, J.L. Spreading depolarization in the brainstem mediates sudden cardiorespiratory arrest in mouse SUDEP models. Science Translational Medicine 7, 9 (2015).

28. Stern, M.A., Cole, E.R., Gross, R.E. & Berglund, K. Seizure event detection using intravital two-photon calcium imaging data. Neurophotonics 11, 024202 (2024).

29. Basarsky, T.A., Duffy, S.N., Andrew, R.D. & MacVicar, B.A. Imaging spreading depression and associated intracellular calcium waves in brain slices. Journal of Neuroscience 18, 7189–7199 (1998).

30. Nedergaard, M., Cooper, A.J.L. & Goldman, S.A. Gap-Junctions are Required for the Propagation of Spreading Depression. Journal of Neurobiology 28, 433–444 (1995).

31. Enger, R., et al. Dynamics of Ionic Shifts in Cortical Spreading Depression. Cerebral Cortex 25, 4469–4476 (2015).

32. Lado, F.A. & Moshé, S.L. How do seizures stop? Epilepsia 49, 1651–1664 (2008).

33. Inoue, M., et al. Rational Engineering of XCaMPs, a Multicolor GECI Suite for In Vivo Imaging of Complex Brain Circuit Dynamics. Cell 177, 1346–1360 (2019).

34. Deng, X.N., et al. Tuning Protein Dynamics to Sense Rapid Endoplasmic-Reticulum Calcium Dynamics. Angewandte Chemie-International Edition 60, 23289–23298 (2021).

35. André, V., Pineau, N., Motte, J.E., Marescaux, C. & Nehlig, A. Mapping of neuronal networks underlying generalized seizures induced by increasing doses of pentylenetetrazol in the immature and adult rat:: a c-Fos immunohistochemical study. European Journal of Neuroscience 10, 2094–2106 (1998).

36. Nathanson, J.L., Yanagawa, Y., Obata, K. & Callaway, E.M. Preferential labeling of inhibitory and excitatory cortical neurons by endogenous tropism of adeno-associated virus and lentivirus vectors. Neuroscience 161, 441–450 (2009).

37. Liou, J.Y., et al. Multivariate regression methods for estimating velocity of ictal discharges from human microelectrode recordings. Journal of Neural Engineering 14, 11 (2017).

38. Smith, E.H., et al. Human interictal epileptiform discharges are bidirectional traveling waves echoing ictal discharges. Elife 11, 20 (2022).

39. Marshall, W.H. Spreading cortical depression of Leao. Physiological Reviews 39, 239–279 (1959).

40. Heck, C.N., et al. Two-year seizure reduction in adults with medically intractable partial onset epilepsy treated with responsive neurostimulation: Final results of the RNS System Pivotal trial. Epilepsia 55, 432–441 (2014).

41. Geller, E.B., et al. Brain-responsive neurostimulation in patients with medically intractable mesial temporal lobe epilepsy. Epilepsia 58, 994–1004 (2017).

42. Hogan, P.G., Lewis, R.S. & Rao, A. Molecular Basis of Calcium Signaling in Lymphocytes: STIM and ORAI. in Annual Review of Immunology, Vol 28 (ed. W.E. Paul, D.R. Littman & W.M. Yokoyama) 491–533 (Annual Reviews, Palo Alto, 2010).

43. Gilbert, G., et al. Calcium Signaling in Cardiomyocyte Function. Cold Spring Harbor Perspectives in Biology 12, 29 (2020).

44. Akerboom, J., et al. Crystal Structures of the GCaMP Calcium Sensor Reveal the Mechanism of Fluorescence Signal Change and Aid Rational Design. Journal of Biological Chemistry 284, 6455–6464 (2009).

45. Ali, F. & Kwan, A.C. Interpreting in vivo calcium signals from neuronal cell bodies, axons, and dendrites: a review. Neurophotonics 7, 17 (2020).

46. Kucharz, K. & Lauritzen, M. CaMKII-dependent endoplasmic reticulum fission by whisker stimulation and during cortical spreading depolarization. Brain 141, 1049–1062 (2018).

47. Khoshkhoo, S., Vogt, D. & Sohal, V.S. Dynamic, Cell-Type-Specific Roles for GABAergic Interneurons in a Mouse Model of Optogenetically Inducible Seizures. Neuron 93, 291–298 (2017).

48. Tran, C.H., et al. Interneuron Desynchronization Precedes Seizures in a Mouse Model of Dravet Syndrome. Journal of Neuroscience 40, 2764–2775 (2020).

49. Farrell, J.S., et al. In vivo assessment of mechanisms underlying the neurovascular basis of postictal amnesia. Scientific Reports 10, 13 (2020).

50. Heuser, K., et al. Ca2+ Signals in Astrocytes Facilitate Spread of Epileptiform Activity. Cerebral Cortex 28, 4036–4048 (2018).

51. Van Harreveld, A. & Stamm, J.S. Consequences of cortical convulsive activity in rabbit. Journal of Neurophysiology 17, 505–520 (1954).

52. Van Harreveld, A. & Stamm, J.S. Cortical response to metrazol and sensory stimulation in the rabbit. Electroencephalography and Clinical Neurophysiology 7, 363–370 (1955).

53. Vergnes, M. & Marescaux, C. Cortical and thalamic lesions in rats with genetic absence epilepsy. Journal of Neural Transmission-General Section, 71–83 (1992).

54. Vinogradova, L., Kuznetsova, G. & Coenen, A. Audiogenic seizures associated with a cortical spreading depression wave suppress spike-wave discharges in rats. Physiology & Behavior 86, 554–558 (2005).

55. Somjen, G.G. Mechanisms of spreading depression and hypoxic spreading depression-like depolarization. Physiological Reviews 81, 1065–1096 (2001).

56. Wei, Y., Ullah, G. & Schiff, S. Unification of Neuronal Spikes, Seizures, and Spreading Depression. Journal of Neuroscience 34, 11733–11743 (2014).

57. Dietz, R.M., Weiss, J.H. & Shuttleworth, C.W. Zn2+ influx is critical for some forms of spreading depression in brain slices. Journal of Neuroscience 28, 8014–8024 (2008).

58. Pal, S., Sombati, S., Limbrick, D.D. & DeLorenzo, R.J. In vitro status epilepticus causes sustained elevation of intracellular calcium levels in hippocampal neurons. Brain Research 851, 20–31 (1999).

59. Sham, J.S.K., Cleemann, L. & Morad, M. Functional coupling of Ca2+ channels and ryanodine receptors in cardiac myocytes. Proceedings of the National Academy of Sciences of the United States of America 92, 121–125 (1995).

60. Furuichi, T., et al. Multiple types of ryanodine receptor Ca2+ release channels are differentially expressed in rabbit brain. Journal of Neuroscience 14, 4794–4805 (1994).

61. Lanner, J.T., Georgiou, D.K., Joshi, A.D. & Hamilton, S.L. Ryanodine Receptors: Structure, Expression, Molecular Details, and Function in Calcium Release. Cold Spring Harbor Perspectives in Biology 2, 21 (2010).

62. Copello, J.A., Barg, S., Onoue, H. & Fleischer, S. Heterogeneity of Ca2+ gating of skeletal muscle and cardiac ryanodine receptors. Biophysical Journal 73, 141–156 (1997).

63. Fill, M. & Copello, J.A. Ryanodine receptor calcium release channels. Physiological Reviews 82, 893–922 (2002).

64. Foskett, J.K., White, C., Cheung, K.H. & Mak, D.O.D. Inositol trisphosphate receptor Ca2+ release channels. Physiological Reviews 87, 593–658 (2007).

65. Giorgi, C., Marchi, S. & Pinton, P. The machineries, regulation and cellular functions of mitochondrial calcium. Nature Reviews Molecular Cell Biology 19, 713–730 (2018).

66. Zhang, S.Y.L., et al. STIM1 is a Ca2+ sensor that activates CRAC channels and migrates from the Ca2+ store to the plasma membrane. Nature 437, 902–905 (2005).

67. Park, C.Y., et al. STIM1 Clusters and Activates CRAC Channels via Direct Binding of a Cytosolic Domain to Orai1. Cell 136, 876–890 (2009).

68. Pietrobon, D. & Moskowitz, M.A. Chaos and commotion in the wake of cortical spreading depression and spreading depolarizations. Nature Reviews Neuroscience 15, 379–393 (2014).

69. Aiba, I., Wehrens, X.H.T. & Noebels, J.L. Leaky RyR2 channels unleash a brainstem spreading depolarization mechanism of sudden cardiac death. Proceedings of the National Academy of Sciences of the United States of America 113, E4895–E4903 (2016).

70. Wuytack, F., Raeymaekers, L. & Missiaen, L. Molecular physiology of the SERCA and SPCA pumps. Cell Calcium 32, 279–305 (2002).

71. Sato, S.S., et al. Simultaneous two-photon imaging of intracellular chloride concentration and pH in mouse pyramidal neurons in vivo. Proceedings of the National Academy of Sciences of the United States of America 114, E8770–E8779 (2017).

72. Sun, X.L., et al. Simultaneous monitoring of intracellular pH changes and hemodynamic response during cortical spreading depression by fluorescence-corrected multimodal optical imaging. Neuroimage 57, 873–884 (2011).

73. Menyhárt, A., et al. Age or ischemia uncouples the blood flow response, tissue acidosis, and direct current potential signature of spreading depolarization in the rat brain. American Journal of Physiology-Heart and Circulatory Physiology 313, H328–H337 (2017).

74. Menyhárt, A., et al. Spreading depolarization remarkably exacerbates ischemia-induced tissue acidosis in the young and aged rat brain. Scientific Reports 7, 13 (2017).

75. Kim, J.H., et al. Noninvasive measurement of the pH of the endoplasmic reticulum at rest and during calcium release. Proceedings of the National Academy of Sciences of the United States of America 95, 2997–3002 (1998).

76. Dreier, J.P., et al. Terminal spreading depolarization and electrical silence in death of human cerebral cortex. Annals of Neurology 83, 295–310 (2018).

77. Carlson, A.P., et al. Terminal spreading depolarizations causing electrocortical silencing prior to clinical brain death: case report. Journal of Neurosurgery 131, 1773–1779 (2019).

78. Hernandez-Caceres, J., Macias-Gonzalez, R., Brozek, G. & Bures, J. Systemic ketamine blocks cortical spreading depression but does not delay the onset of terminal anoxia depolarization in rats. Brain Research 437, 360–364 (1987).

79. Müller, M. & Somjen, G.G. Inhibition of major cationic inward currents prevents spreading depression-like hypoxic depolarization in rat hippocampal tissue slices. Brain Research 812, 1–13 (1998).

80. Collins, F., Schmidt, M.F., Guthrie, P.B. & Kater, S.B. Sustained increase in intracellular calcium promotes neuronal survival. Journal of Neuroscience 11, 2582–2587 (1991).

81. Franklin, J.L. & Johnson, E.M. Suppression of programmed neuronal death by sustained elevation of cytoplasmic calcium. Trends in Neurosciences 15, 501–508 (1992).

82. Heinemann, U. & Lux, H.D. Ceiling of stimulus induced rises in extracellular potassium concentration in the cerebral-cortex of cat. Brain Research 120, 231–249 (1977).

83. Anderson, D., et al. Closed-loop stimulation in periods with less epileptiform activity drives improved epilepsy outcomes. Brain 147, 521–531 (2024).

84. Huang, X.P., et al. AAV2 production with optimized N/P ratio and PEI-mediated transfection results in low toxicity and high titer for in vitro and in vivo applications. J. Virol. Methods 193, 270–277 (2013).

85. Goldey, G.J., et al. Removable cranial windows for long-term imaging in awake mice. Nature Protocols 9, 2515–2538 (2014).

86. Roome, C.J. & Kuhn, B. Chronic cranial window with access port for repeated cellular manipulations, drug application, and electrophysiology. Frontiers in Cellular Neuroscience 8, 8 (2014).

87. Roome, C.J. & Kuhn, B. Voltage Imaging with ANNINE Dyes and Two-Photon Microscopy. in Multiphoton Microscopy (ed. E. Hartveit) 297–334 (Humana Press Inc, Totowa, 2019).

88. Moucka, R., Sedlacík, M., Osicka, J. & Pata, V. Mechanical properties of bulk Sylgard 184 and its extension with silicone oil. Scientific Reports 11, 9 (2021).

89. Stern, M.A., et al. Applications of Bioluminescence-Optogenetics in Rodent Models. Methods Mol Biol 2525, 347–363 (2022).

90. Jiang, S., et al. Automated, highly reproducible, wide-field, light-based cortical mapping method using a commercial stereo microscope and its applications. Biomed. Opt. Express 7, 3478–3490 (2016).

91. Ferezou, I., et al. Spatiotemporal dynamics of cortical sensorimotor integration in behaving mice. Neuron 56, 907–923 (2007).

92. Mason, C.R. & Cooper, R.M. Permanent change in convulsive threshold in normal and brain-damaged rats with repeated small doses of pentylenetetrazol. Epilepsia 13, 663-& (1972).

93. Schindelin, J., et al. Fiji: an open-source platform for biological-image analysis. Nature Methods 9, 676–682 (2012).

94. Pachitariu, M., et al. Suite2p: beyond 10,000 neurons with standard two-photon microscopy. bioRxiv, 061507 (2017).

95. Stringer, C., Wang, T., Michaelos, M. & Pachitariu, M. Cellpose: a generalist algorithm for cellular segmentation. Nature Methods 18, 100–106 (2021).

96. Wenzel, M., Hamm, J.P., Peterka, D.S. & Yuste, R. Reliable and Elastic Propagation of Cortical Seizures In Vivo. Cell Reports 19, 2681–2693 (2017).

97. Somarowthu, A., Goff, K.M. & Goldberg, E.M. Two-photon calcium imaging of seizures in awake, head-fixed mice. Cell Calcium 96, 8 (2021).

98. Rolston, J.D., Laxpati, N.G., Gutekunst, C.-A., Potter, S.M. & Gross, R.E. Spontaneous and evoked high-frequency oscillations in the tetanus toxin model of epilepsy. Epilepsia 51, 2289–2296 (2010).

