## Supplemental Material for "Organellular imaging *in vivo* reveals a depletion of endoplasmic reticular calcium during post-ictal cortical spreading depolarization"

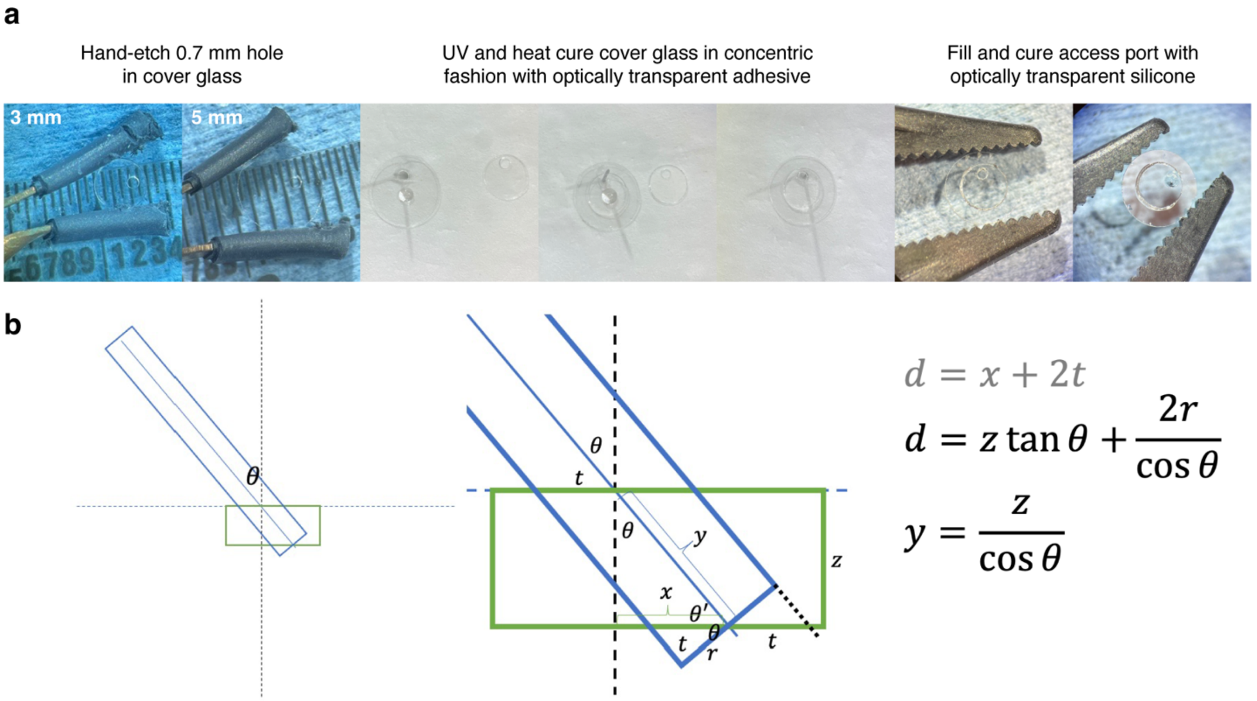


**Supplementary Fig. 1. Cranial window with access port construction**

**(a)** Glass coverslips (3 and 5mm, #1 thickness) are individually hand etched using a fine stone conical burr to generate 0.5-0.7mm holes. These are then affixed to one another concentrically using optically transparent adhesive in a staged process, where the holes are aligned by threading the glass onto an erected micro dissection needle (000) and then cured using UV light and heat. These holes are then filled with optically transparent silicone and cured at room temperature under vacuum and then with heat to optimize the strength and elasticity of the silicone. **(b)** Schematic of pipette approach (blue) to horizontal concentric window access port (green) with relevant equations to calculate the minimum access port size (d), and stereotaxic-z (y) needed to traverse the window given a pipette radius (r), an angle of approach to the vertical (θ), and window thickness (z). For example, a recording pipette with a radius of 50 µm at an approach angle of 40$^{o}$ for a 3-layer concentric window of #1 cover glass (150 µm/layer) would need an access port of at least 508µm and would need 587µm in stereotactic-z to traverse the window.

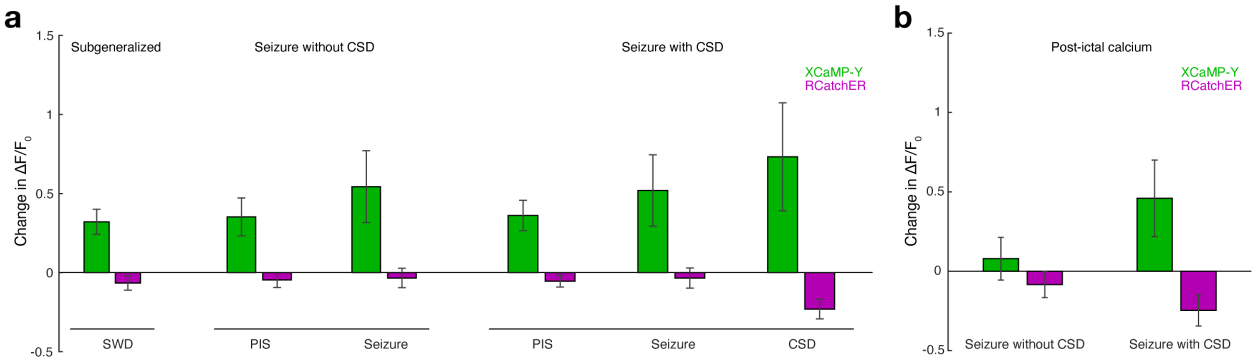


**Supplementary Fig. 2. Calcium changes within subject for representative subject across sub-generalized recordings and seizures with and without CSD**

**(a)** Calcium levels (mean change with standard error) during each event (PIS, seizure and CSD) stratified by recording type for the individual subject presented in figure 2D. **(b)** Post-ictal calcium change comparing the seizure recording with and without CSD (mean change with standard error).

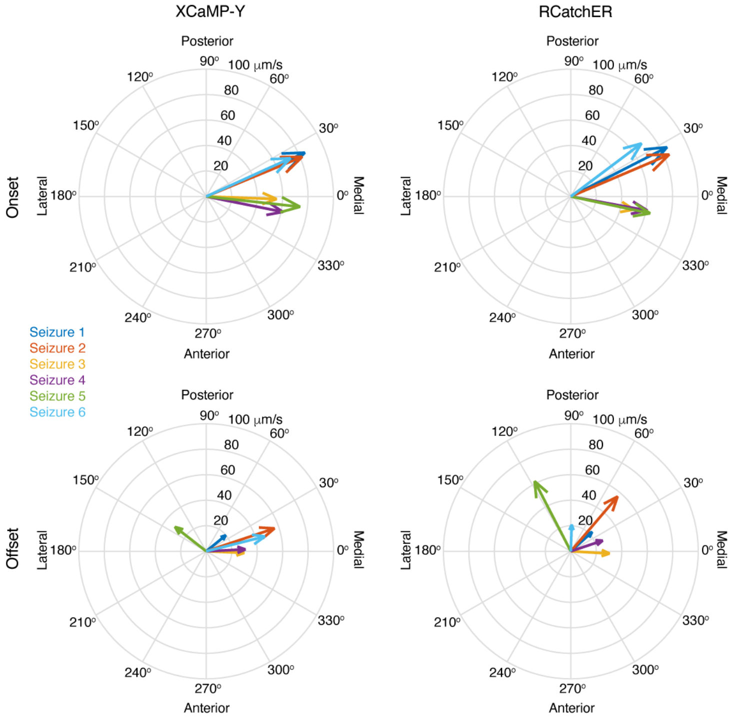


**Supplementary Fig. 3. Polar plots of all CSDs that occurred with seizure**

Polar plot of the CSD onset and offset propagation vectors modeled by applying spatial linear regression to the neuronal recruitment times across all seizures, showing wavefront direction (vector angle) and propagation speed (vector magnitude). Each CSD is depicted using a unique color matched to Fig. 3.

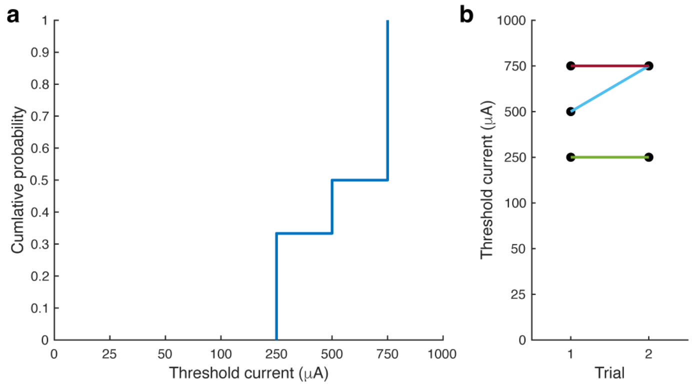


**Supplementary Fig. 4. Threshold current to induce CSD with RNS waveform parameters**

**(a)** Cumulative probability of inducing a CSD across increasing stimulation current (N=6 recordings across 3 subjects). **(b)** Threshold current repeated across trials within subject.

**Legends for Movies**

**Supplementary Movie 1. Representative generalized seizure with post-ictal CSD**
In vivo awake two-photon imaging of cytosolic (XCaMP-Y; green) and ER luminal (RCatchER; magenta) calcium during the progression of a PTZ-induced generalized seizure from the pre-ictal through the post-ictal phase. Field of view is 450mm×450mm. Playback speed at 5x. (MPEG-4, 51.7MB).

**Supplementary Movie 2. Representative electrical stimulation-induced CSD**
In vivo awake two-photon imaging of cytosolic (XCaMP-Y; green) and ER luminal (RCatchER; magenta) calcium during the progression of an electrically stimulated CSD. Field of view is 450mm×450mm. Playback speed at 5x. (MPEG-4, 52.1MB).
